# Genome-wide ribosome profiling uncovers the role of iron in the control of protein translation

**DOI:** 10.1101/2022.09.22.509115

**Authors:** Hanna Barlit, Antonia M. Romero, Ali Gülhan, Praveen K. Patnaik, Alexander Tyshkovskiy, María T. Martínez-Pastor, Vadim N. Gladyshev, Sergi Puig, Vyacheslav M. Labunskyy

## Abstract

Iron is an essential trace element that serves as a cofactor for enzymes involved in multiple metabolic pathways, including ribosome biogenesis, protein translation, DNA synthesis and repair, lipid metabolism, and mitochondrial oxidative phosphorylation. In eukaryotes, iron deficiency leads to global inhibition of protein synthesis and coordinated changes in gene expression to limit iron utilization. Although several steps of protein translation depend on iron-containing enzymes, the contribution of iron to the translation process is not understood at the molecular level. Here, we report a genome-wide analysis of protein translation in response to iron deficiency in yeast using ribosome profiling. We show that iron depletion affects global protein synthesis as well as leads to translational repression of several groups of genes involved in iron-related processes. We further demonstrate that the RNA-binding proteins Cth1 and Cth2 play a central role in controlling the changes in protein translation by repressing the activity of the iron-dependent Rli1 ribosome recycling factor, inhibiting mitochondrial translation, and affecting the translation of genes involved in heme biosynthesis. We also discovered a mechanism, whereby iron deficiency represses translation of *MRS3* mRNA, encoding mitochondrial iron transporter, through increased expression of antisense long non-coding RNA. Together, our results reveal complex gene expression and protein synthesis remodeling in response to low iron showing how this important metal affects protein translation at multiple levels.

## Introduction

Iron (Fe) is an essential trace element that serves as a cofactor for enzymes involved in numerous cellular processes, including protein translation, DNA replication and repair, lipid biosynthesis, and mitochondrial oxidative phosphorylation (Zhang, 2014). Dysregulation of Fe homeostasis has been implicated in health and disease. In humans, Fe deficiency causes anemia and leads to the decline of immune, neuronal, and muscle functions (Andrews, 2000; Cronin et al., 2019). On the other hand, Fe overload leads to organ failure, due to Fe toxicity, and the development of several disorders, such as Alzheimer’s disease, cancer, and delayed wound healing (Hare et al., 2013). Moreover, imbalance in Fe metabolism has been implicated in aging and age-associated organ failure, but the exact mechanisms remain unclear (Xu et al., 2012). A better understanding of the mechanisms by which cells adapt to changes in Fe availability is required for the development of targeted therapeutic strategies against diseases associated with the dysregulation of Fe homeostasis.

Yeast *Saccharomyces cerevisiae* has proven to be a useful model for understanding the basic mechanisms of adaptation of eukaryotic cells to Fe depletion. Previous studies in yeast have shown that Fe deficiency leads to significant changes in gene expression (Puig et al., 2005; Romero et al., 2019; Shakoury-Elizeh et al., 2004). These changes are mediated by the Aft1 and Aft2 transcription factors, which activate the transcription of ∼35 genes known as the “Fe regulon”. Expression of these genes allows cellular adaptation to low Fe through several mechanisms: (i) increasing Fe uptake into the cell, (ii) mobilizing Fe from vacuolar stores, and (iii) recycling iron by promoting heme degradation (Philpott and Protchenko, 2008; Ramos-Alonso et al., 2020; Shakoury-Elizeh *et al*., 2004). In addition, cells adapt to Fe deficiency by dropping the utilization of Fe for non-essential metabolic pathways and shifting to Fe-independent metabolism through the increased expression of the mRNA-binding protein Cth2, which is under the control of the Aft1/2 transcription factors. Cth2 recognizes and binds to AU-rich sequences (5’-UAUUUAUU-3’ and 5’-UUAUUUAU-3’) in the 3’-untranslated region (UTR) of several mRNAs coding for proteins containing Fe as a cofactor (including aconitase, succinate dehydrogenase, and components of the mitochondrial electron transport chain) and stimulate their degradation (Puig *et al*., 2005). A paralog of Cth2, named Cth1, is 46% identical to Cth2, recognizes similar consensus sequences, and can repress the expression of a partially overlapping set of target genes (Puig *et al*., 2005; Puig et al., 2008; Ramos-Alonso et al., 2018). Thus, Cth1 and Cth2 remodel Fe metabolism prioritizing Fe for proteins involved in essential processes such as DNA replication and repair (Sanvisens et al., 2011).

While transcriptional responses to Fe deficiency have been extensively characterized, little is known about the role of Fe in regulating protein translation. Previous reports have shown that Fe deficiency leads to a global inhibition of protein synthesis, which is dependent on the TORC1 and Gcn2/eIF2α pathways (Romero et al., 2020; Romero *et al*., 2019). In addition, Fe serves as a cofactor for several enzymes participating in protein translation, including proteins involved in modifications of factors affecting translation elongation, post-transcriptional transfer RNA (tRNA) modifications, translational termination, and ribosome recycling (Keeling et al., 2006; Romero et al., 2021; Young et al., 2015). Moreover, Cth2 has been shown to repress the translation of specific transcripts in response to Fe deficiency (Ramos-Alonso *et al*., 2018). However, changes in protein translation at the genome-wide level and the mechanisms of translational regulation by Fe have not been previously investigated.

In this work, we applied RNA-Seq and ribosome profiling (Ribo-Seq) to yeast cells undergoing adaptation to Fe deficiency to quantitatively measure translational changes genome-wide. At the translation level, we uncovered that Fe depletion leads to specific downregulation of genes involved in Fe-dependent processes, including mitochondrial translation and heme biosynthesis. We further show that this translational regulation is mediated by Cth1 and Cth2. We find that Fe deficiency also affects global protein translation by dramatically decreasing the activity of the ribosome recycling factor Rli1. Conversely, eliminating Cth1 and Cth2 increased the levels of Rli1 indicating that cells adapt to Fe deficiency by limiting the activity of Rli1 in response to low Fe in a Cth1/2-dependent manner. Our data also demonstrate that Fe deficiency affects transcription of antisense long non-coding RNAs (lncRNAs) that play a regulatory role by repressing cognate sense mRNA translation. Taken together, this study uncovers the role of the RNA-binding proteins Cth1 and Cth2 in the control of protein translation and how Fe regulates protein synthesis at both global and transcript-specific levels.

## Results

### Transcriptional and translational responses during the adaptation to iron deficiency

Previous studies have shown that Fe deficiency in yeast *S. cerevisiae* leads to changes in transcriptional profiles (Hausmann et al., 2008; Puig *et al*., 2005; Shakoury-Elizeh *et al*., 2004). In addition, prolonged Fe deficiency has been shown to induce global inhibition of protein synthesis, which is dependent on the Gcn2/eIF2α pathway (Romero *et al*., 2020). However, little is known about how Fe deprivation manifests at the level of translational regulation. To distinguish the effects of Fe deficiency on mRNA abundance from its effects on protein translation, we performed RNA-Seq and ribosome profiling (Ribo-Seq) in wild-type W303a yeast cells cultured in Fe-sufficient (+Fe) or Fe-deficient conditions (-Fe), achieved by the addition of the Fe^2+^-specific chelator bathophenanthroline disulfonate (BPS). For this, exponentially growing cells were isolated under Fe-sufficient conditions, and exposed to a short-(3 h) or long-term (6 h) Fe deficiency (**Fig 1A, Dataset S1**). In addition to wild-type cells, we monitored genome-wide transcriptional and translational changes upon Fe limitation in the *cth1*Δ*cth2*Δ mutant. The RNA-Seq and Ribo-Seq profiles showed significantly altered gene expression patterns during adaptation to low Fe from 0 h to 6 h of Fe deficiency, with intermediate changes at 3 h (**Fig S1**). Consistent with previous studies, we also observed strong inhibition of global protein synthesis after 6 h of Fe deficiency in both wild-type cells and the *cth1*Δ*cth2*Δ mutant as measured by polysome analysis, but changes at 3 h were less pronounced (**Fig 1B**). We found that most of the changes in translation (Ribo-Seq) during Fe deficiency correlated with the changes in mRNA abundance (RNA-Seq) (**Fig 1C**). These data demonstrate that the responses to Fe deficiency are mediated primarily by changes in mRNA levels. However, we also found a group of genes that were altered specifically at the translation level indicating that an additional level of regulation contributes to the response of yeast cells to low Fe (**Fig 1D**).

**Fig 1.**
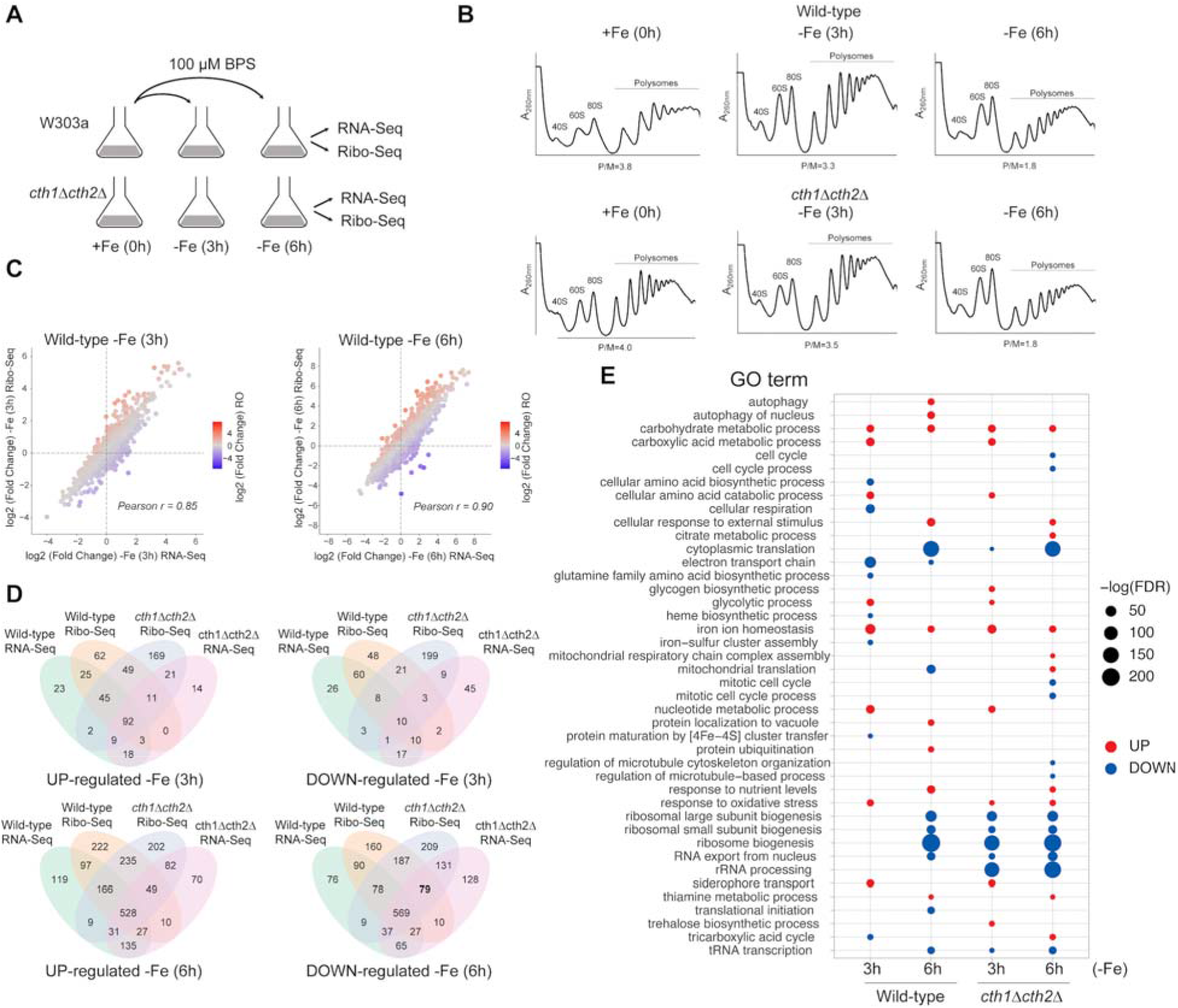
Coordinated changes in mRNA levels and translation allow metabolic reprogramming in response to low Fe. **A**) Experimental design. **B**) Polysome profiles of wild-type and *cth1*Δ*cth2*Δ cells during the time-course of Fe deficiency. Polysome to monosome (P/M) ratios were calculated using areas under the curve using ImageJ. Corresponding monosome (40S, 60S, 80S) and polysome peaks are indicated. **C**) Comparison of transcriptional (RNA-Seq) and translational (Ribo-Seq) changes during the course of Fe deficiency. Genes whose ribosome occupancy (RO) is increased (red) or decreased (blue) during Fe deficiency are highlighted. **D**) Venn diagrams show genes whose transcription (RNA-Seq) and translation (Ribo-Seq) were significantly changed during Fe deficiency (FDR<0.05). **E**) GO terms enriched among translationally up- and down-regulated genes during Fe deficiency.

Our analysis of the RNA-Seq and Ribo-Seq data revealed many of the previously known changes in response to Fe deficiency. Among the genes transcriptionally up-regulated during the time-course of Fe deficiency, we observed a number of Fe regulon genes that are controlled by the Aft1 and Aft2 transcription factors, which validated our experimental approach (Philpott et al., 2012; Ramos-Alonso *et al*., 2020). Genes up-regulated in response to Fe limitation were enriched in GO categories “iron ion homeostasis”, “siderophore transport”, “carbohydrate metabolic process”, “glycolytic process”, “nucleotide metabolic process”, and “cellular amino acid catabolic process” (**Fig 1E**, **Dataset S2**). For example, expression of genes involved in Fe siderophore uptake (*FIT1-3, ARN1-4*), reductive Fe uptake (*FRE1-FRE4, FET3, FTR1, ATX1, CCC2, FET4*), and mobilization of vacuolar Fe (*FRE6, FET5, FTH1, SMF3*) increased several folds in response to Fe deprivation in wild-type cells (**Fig S2**). We also observed that expression levels of *CTH2*, a gene coding for the mRNA-binding protein involved in metabolic remodeling in response to low Fe, were significantly induced in wild-type cells during Fe deficiency.

Among downregulated genes in response to Fe limitation, the most enriched GO categories included “cytoplasmic translation”, “ribosomal large and small subunits”, and “ribosome biogenesis” consistent with the overall inhibition of protein synthesis. However, there were substantial differences in downregulated genes between wild-type and *cth1*Δ*cth2*Δ cells. For example, “cellular respiration”, “electron transport chain”, “heme biosynthetic process”, “iron-sulfur cluster assembly”, “mitochondrial translation”, and “tricarboxylic acid cycle” were enriched among downregulated genes in wild-type cells, but not in the *cth1*Δ*cth2*Δ mutant. In contrast, “rRNA processing” and “mitotic cell cycle” were specifically downregulated in the *cth1*Δ*cth2*Δ cells suggesting a role of Cth1 and Cth2 in regulation of these processes.

### Ribo-Seq analyses reveal the role of the RNA-binding proteins Cth1 and Cth2 in translational regulation

During Fe deficiency, yeast cells activate the expression of Cth2, an mRNA-binding protein involved in the remodeling of Fe metabolism. In response to low Fe, Cth2 and, to a lower extent, Cth1 post-transcriptionally inhibit the expression of several mRNAs that contain AU-rich elements (AREs) in the 3’-UTR (Puig *et al*., 2005; Ramos-Alonso *et al*., 2018). We expected that the lack of Cth1 and Cth2 in the *cth1*Δ*cth2*Δ mutant would lead to the derepression of their targets. By comparing changes in protein synthesis profiles (Ribo-Seq) in wild-type and *cth1*Δ*cth2*Δ cells we identified 164 genes that were translationally up-regulated in the *cth1*Δ*cth2*Δ mutant compared to wild-type cells during 6 h of Fe deficiency (FDR<0.05) (**Fig 2A** and **Table S1**). Among these translationally up-regulated genes, we found several previously reported targets of Cth1 and Cth2 that code for components of the electron transport chain (ETC) and tricarboxylic acid (TCA) cycle (**Fig 2B** and **Fig S3**) consistent with the known function of Cth1/2 mRNA-binding proteins in post-transcriptional regulation of Fe-dependent genes (Puig *et al*., 2008; Ramos-Alonso *et al*., 2018). In addition to previously described Cth1/2 targets, our data uncovered that Cth1 and Cth2 are limiting the expression of genes encoding for mitochondrial ribosomal proteins (MRPs) (36 ribosomal proteins of the large subunit and 24 ribosomal proteins of the small subunit) as well as proteins involved in the assembly of the ETC (16 genes), mitochondrial translation (15 genes), protein translocation to mitochondria (10 genes), ATP synthase assembly (4 genes), and heme biosynthesis (4 genes) (**Fig 2C**). Because known targets of Cth1 and Cth2 contain AREs in their 3’UTR, we searched for putative AREs within 3’UTR of these genes (**Table S1**). This analysis revealed several transcripts that are directly regulated by Cth1/2. For example, we identified mRNA transcripts encoding Cbp3-Cbp4-Cbp6 complex, which is involved in the translation and assembly of the mitochondrial cytochrome bc1 complex (Gruschke et al., 2012). Our data demonstrate that *CBP3, CBP4,* and *CBP6* are coordinately downregulated in response to low Fe in wild-type cells at the mRNA level (but not in the *cth1*Δ*cth2*Δ mutant), suggesting that *CBP3, CBP4,* and *CBP6* are directly regulated by Cth1/2 during Fe deficiency (**Fig 2D**). In contrast, most of the genes encoding for MRP proteins lack AREs in their 3’UTRs and Fe deficiency decreased their translation without affecting their transcript levels (**Fig 2E** and **Fig S4**). We also observed that expression of Cth1 and Cth2 is reversely correlated with the synthesis of mitochondrial ribosomal proteins of the small subunit (MRPS) and large subunit (MRPL) during the shift of cells from the fermentable carbon source to the non-fermentable medium containing glycerol (**Fig S5**) that requires the induction of the mitochondrial ETC components and mitochondrial translation (Isaac et al., 2018). Further analysis of MRP genes activated by Fe deficiency in the *cth1*Δ*cth2*Δ mutant identified the presence of Puf3 binding sites in their 3’UTR sequences (**Table S1**). Puf3 is an RNA-binding protein that has been implicated in the control of mitochondrial translation (Couvillion et al., 2016). In yeast, Puf3 senses glucose availability and binds to the 3’UTRs of mRNA transcripts involved in mitochondrial translation, including genes encoding components of the mitochondrial large and small ribosomal protein subunits, promoting their degradation. However, upon glucose depletion, Puf3 becomes phosphorylated and switches its function to promote the translation of its target mRNAs (Lee and Tu, 2015). Thus, increased translation of MRPs in the *cth1*Δ*cth2*Δ mutant could be mediated by an increased activity of the Puf3 mRNA-binding protein.

**Fig 2.**
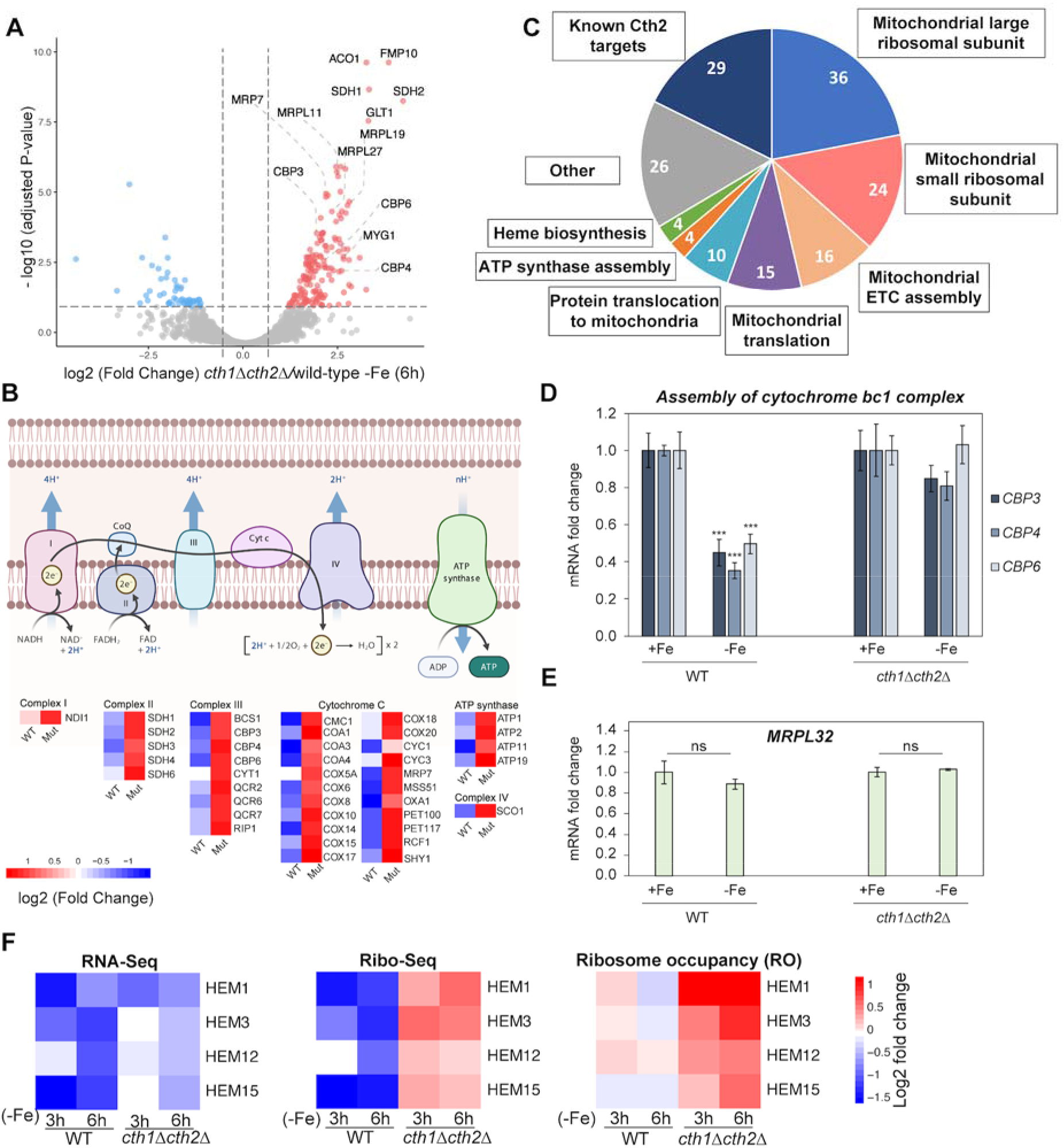
Fe deficiency leads to Cth1/2-dependent inhibition of mitochondrial translation and heme biosynthesis. **A**) Volcano plot of differentially regulated genes in the *cth1*Δ*cth2*Δ mutant compared to wild-type cells at 6 h (-Fe). **B**) Heatmap of differentially regulated genes of ETC in *cth1*Δ*cth2*Δ compared to wild-type cells during 6 h of Fe deficiency (FDR<0.05). **C**) Pie chart of genes activated in the *cth1*Δ*cth2*Δ compared to wild-type cells. **D)** Fe deficiency coordinately downregulates expression of transcripts encoding the Cbp3-Cbp4-Cbp6 complex. The expression of *CBP3, CBP4,* and *CBP6* was determined by RT-qPCR. Error bars represent SEM of three independent experiments, ***p<0.001 (one-way ANOVA). **E)** Translation of mitochondrial ribosomal proteins is regulated by Fe deficiency in a Cth1/2-dependent manner without affecting mRNA transcript levels. The expression of *MRPL32* was determined by RT-qPCR. Error bar represent SEM of three independent experiments, ns, non-significant (one-way ANOVA). **F)** Heme biosynthesis is translationally regulated by Cth1 and Cth2. Heatmaps show log2 fold change in mRNA abundance (RNA-Seq), protein translation (Ribo-Seq), and ribosome occupancy (RO) compared to untreated (0 h) samples.

Notably, we found that Fe deficiency also leads to translational downregulation of genes encoding enzymes involved in heme biosynthesis, including *HEM1, HEM3, HEM12, HEM15* (**Fig 2F**). To estimate translation efficiency and investigate the contribution of transcriptional and translational regulation to gene expression changes, we calculated ribosome occupancy (RO) for each of these genes. In wild-type cells, both mRNA abundance (RNA-seq) and protein synthesis (Ribo-Seq) of this set of genes were significantly decreased by Fe deficiency (FDR<0.05). In contrast, translation of *HEM* transcripts was up-regulated in the *cth1*Δ*cth2*Δ mutant leading to pronounced increase in RO. Together, our results indicate that Cth1 and Cth2 play a central role in controlling the changes in protein translation in response to low Fe by directly and indirectly downregulating multiple transcripts required for mitochondrial translation and heme biosynthesis.

### Fe deficiency leads to increased ribosome occupancy in the 3’UTRs of genes due to decreased activity of Rli1

Although overall protein synthesis is significantly reduced during Fe deficiency, how different steps of protein translation are affected by low Fe is not completely understood. Among enzymes participating in protein translation that require Fe as a cofactor, we selected Rli1 for further analysis. Rli1 is a conserved protein, which requires an iron-sulfur (Fe-S) cluster for its activity and is essential for translation termination and ribosome recycling (Romero *et al*., 2021). *RLI1* mRNA contains two putative AREs within its 3’UTR at 280 and 291 nt from its stop codon, and its transcript levels are up-regulated in cells lacking *CTH1* and *CTH2* as compared to wild-type cells in -Fe conditions (Puig *et al*., 2005; Puig *et al*., 2008) suggesting it is a direct Cth1/2 target mRNA. We expected that, during Fe deficiency, activity of Rli1 would decrease. Consistent with its function in ribosome recycling, we observed diminished Rli1 activity as evidenced by the accumulation of 80S ribosomes in the 3’UTRs of several mRNAs during prolonged Fe deficiency (6 h) in wild-type cells (**Fig 3A**). In contrast, decrease in activity of the Rli1 was delayed in the *cth1*Δ*cth2*Δ mutant. To investigate whether increased ribosome occupancy at 3’UTR in Fe-depleted conditions is associated with active translation, we used yeast strains containing 3xHA tags in the 3’UTRs of *SED1* and *CWP2* (Young *et al*., 2015) downstream of the canonical stop codon. We observed increased levels of 3’UTR translation products for *SED1* and *CWP2* genes during prolonged incubation of wild-type cells in the absence of Fe (**Fig 3B**). Western blot analysis revealed that these bands correspond to the production of small peptide products, rather than read-through translation products that are expected to have larger sizes. Notably, eliminating Cth1 and Cth2 prevented their repressive effects on the expression levels of *RLI1* during Fe deficiency in the *cth1*Δ*cth2*Δ mutant (**Fig 3C**) lowering expression of the *SED1* 3’UTR translation product (**Fig 3D**). Together, these data suggest that Fe deficiency decreases activity of Rli1 by promoting binding of Cth1/2 to the AREs in its 3’UTR and promoting degradation of its transcript.

**Fig 3.**
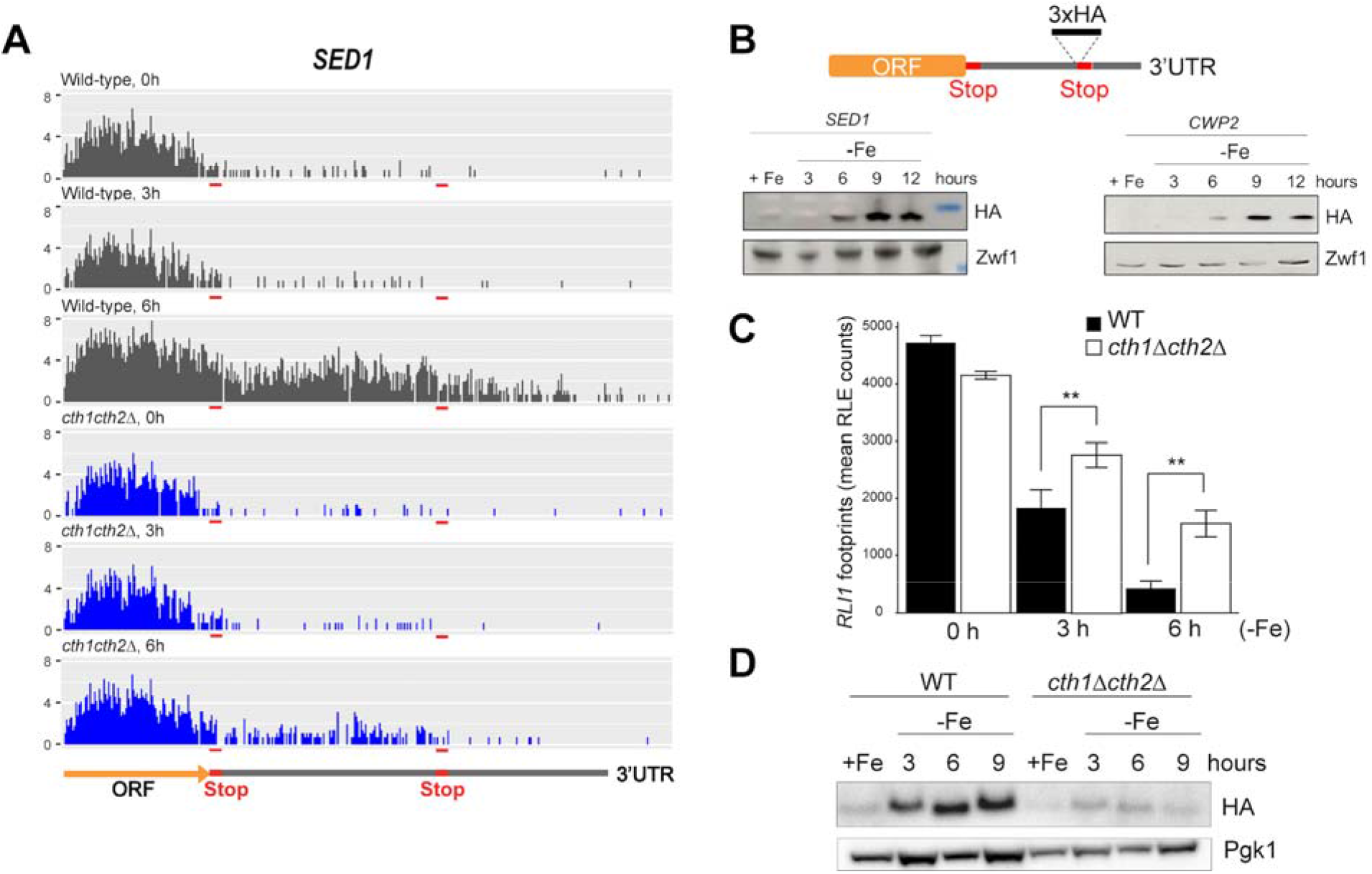
Fe deficiency leads to increased translation of 3’UTRs due to Cth1/2-dependent inhibition of Rli1 activity. **A**) Ribosome occupancy at the 3’UTR of *SED1* mRNA during short (3h) and prolonged (6h) Fe deficiency (-Fe). The location of the translation termination site i shown with red boxes. **B**) Strains containing 3xHA tags inserted in the 3’UTRs of *SED1* and *CWP2* (Young *et al*., 2015) show increased translation downstream of the canonical translation stop codon during prolonged Fe deficiency. Strains expressing tagged proteins were cultured in Fe sufficient (+Fe) media and after Fe deficiency (-Fe) for the indicated time. The expression of tagged proteins was detected by western blot analysis with HA-tag antibody. **C**) Lack of Cth1/2 in the *cth1*Δ*cth2*Δ mutant leads to derepression of *RLI1.* Error bars represent SEM of three independent experiments, **p<0.01 (one-way ANOVA). **D)** Expression of the *SED1* 3’UTR translation product is prevented in the *cth1*Δ*cth2*Δ mutant.

### *MRS3* translation is repressed by antisense transcription of a long non-coding RNA

Among the genes translationally regulated by Fe deficiency, we identified *MRS3* that encodes a mitochondrial Fe transporter. Footprint coverage of *MSR3* was downregulated in response to Fe deficiency suggesting reduced translation of this gene in low Fe conditions (**Fig 4A**, right panel). But at the RNA level, we observed increased expression of the antisense long non-coding RNA (lncRNA), which we named *MRS3^AS^* (**Fig 4A**, left panel). We further confirmed increased expression of the *MRS3^AS^*antisense transcript using RT-qPCR during prolonged Fe deficiency (**Fig 4B**), which was associated with 9.2-fold reduction of the Mrs3 protein levels (**Fig 4C**). In contrast to *MRS3*, low Fe did not affect footprint coverage of its homolog *MRS4,* and we did not observe expression of the antisense RNA for this gene (**Fig S6A**). We then asked whether increased expression of the *MRS3^AS^*transcript is dependent on Cth1 and Cth2. However, we observed increased *MRS3^AS^*transcription in response to low Fe in the *cth1*Δ*cth2*Δ mutant (**Fig S6B**) indicating that Fe deficiency regulates Mrs3 levels in a Cth1/2-independent manner. Finally, to determine if *MRS3^AS^* transcription is sufficient for the repression of *MRS3*, we generated a plasmid expressing antisense lncRNA. Expression of the full length *MRS3^AS^* lncRNA, but not its shorter forms, decreased Mrs3 protein translation in the absence of Fe deficiency (**Fig 4D**) suggesting that ectopic expression of the antisense lncRNA is sufficient to repress protein translation.

**Fig 4.**
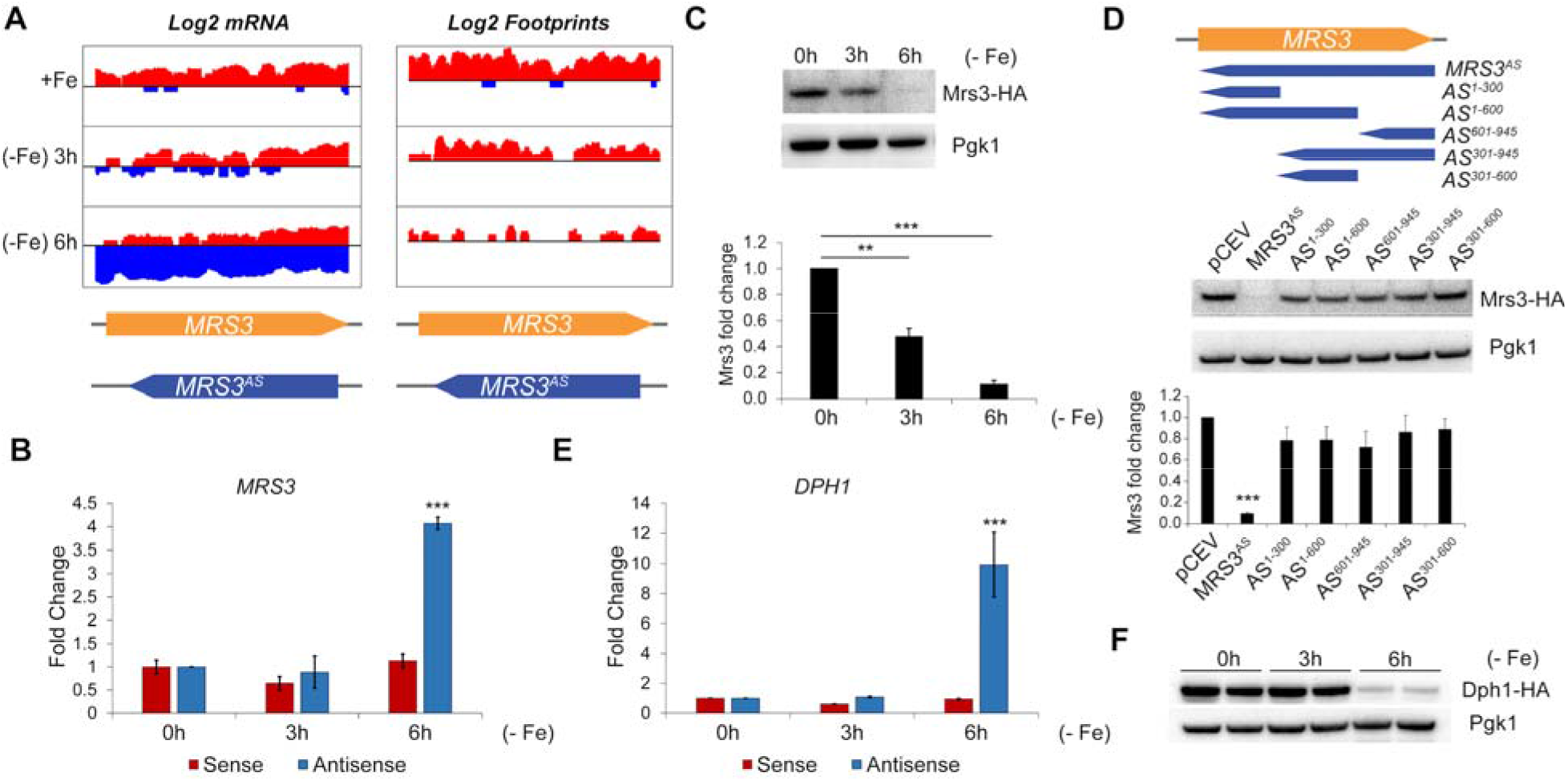
Fe deficiency induces expression of regulatory lncRNAs. **A**) Coverage plot of reads mapped to sense (red) and antisense (blue) *MRS3* transcript. Y-axis represents log2 transformed number of reads, x-axis coordinates show nucleotide positions within *MRS3* ORF. **B**) Expression of antisense *MRS3* (*MRS3*^AS^) transcript is significantly upregulated upon Fe deficiency. Relative expression of *MRS3* and *MRS3*^AS^ transcripts was analyzed by RT-qPCR. Error bars represent SEM of three independent experiments, ***p<0.001 (two-way ANOVA). **C**) Mrs3 protein levels are repressed by prolonged Fe deficiency. Expression of Mrs3-HA and Pgk1 proteins was determined by Western blot with anti-HA and anti-Pgk1 antibodies, respectively. Error bar represent SEM of three independent experiments, **p<0.01, ***p<0.001 (t-test). **D**) Expression of *MRS3^AS^* lncRNA is sufficient to repress translation of *MRS3*. Cells expressing HA-tagged Mrs3 protein were transformed with either an empty vector (pCEV) or pCEV-*MRS3^AS^*expressing full length (*MRS3^AS^*) or short forms of the antisense *MRS3* transcript (*AS*) and Mrs3 protein levels were analyzed by Western blot. Error bars represent SEM of three independent experiments, ***p<0.001 (t-test). **E**) Relative expression of *DPH1* and *DPH1^AS^* transcripts wa analyzed by RT-qPCR. Error bars represent SEM of three independent experiments, ***p<0.001 (two-way ANOVA). **F**) Expression of Dph1-HA protein during Fe deficiency was analyzed by Western blot with anti-HA antibodies.

### Expression of antisense lncRNAs is a conserved regulatory mechanism in yeast

As *MRS3* is translationally regulated by an antisense lncRNA, we asked if other genes might be controlled by low Fe using a similar mechanism. We searched for the presence of antisense lncRNA transcription in other genes that are translationally downregulated by Fe deficiency. For this, we systematically searched for cis-antisense transcripts, which overlap protein coding genes and are transcribed from the opposite DNA strand, leading to decreased sense mRNA translation (**Dataset S3**). In addition to *MRS3,* we identified *DPH1,* which showed increased antisense transcription following 6 h of Fe deficiency (**Fig S6C**), leading to significant decrease of *DPH1* footprint coverage. Using RT-qPCR, we further confirmed the increased expression of the antisense lncRNA (*DPH1^AS^*) during prolonged Fe deprivation (**Fig 4E**), which was associated with decreased translation of the main ORF transcript and decreased levels of Dph1 protein (**Fig 4F**). The observation that translation of both *MRS3* and *DPH1* are repressed by low Fe suggests that this may be a conserved regulatory mechanism.

## Discussion

Although transcriptional responses to Fe deprivation have been extensively characterized, how Fe deficiency affects protein synthesis at the genome-wide level remained elusive. By combining RNA-Seq with the analysis of ribosome occupancy by Ribo-Seq, we quantitatively analyzed translatome changes in response to the short-term and prolonged Fe deficiency in yeast. This allowed us to identify groups of genes, whose expression was altered by low Fe specifically at either transcriptional level or protein synthesis level, and provided mechanistic insights into the role of Fe in protein translation.

The changes in gene expression that we observed were consistent with prior studies that analyzed transcriptomic profiles during Fe deprivation showing altered expression of genes involved in Fe acquisition and mobilization (Puig *et al*., 2005; Shakoury-Elizeh *et al*., 2004). Expression of these genes is activated by the Aft1 and Aft2 transcription factors. In addition, we observed that Cth1 and Cth2 mRNA-binding proteins repress expression of their targets adjusting metabolism to prioritize Fe for essential processes (Philpott *et al*., 2012). For most of the transcripts, a strong correlation between RNA-Seq and Ribo-Seq profiles was found indicating that changes in gene expression induced by Fe deficiency predominantly occur at the mRNA level. However, we also identified a number of genes that were specifically downregulated by Fe deficiency at the translation level. These downregulated proteins included enzymes involved in heme biosynthesis as well as mitochondrial ribosomal proteins and other components of the mitochondrial translational machinery. Translational repression of heme biosynthesis and inhibition of mitochondrial translation were not observed in previous reports because most of the prior studies focused solely on transcriptomic changes.

Importantly, we identified Cth1 and Cth2 as key factors that are responsible for the translational repression. Although the exact mechanisms of translational regulation by Cth1 and Cth2 remain unclear, we found that many of the genes translationally downregulated by Cth1/2 contained binding sites for the Puf3 RNA-binding protein in their 3’UTRs. Puf3 is a known regulator of mitochondrial translation in response to glucose availability, which specifically binds to mRNA and represses translation of mitochondrial ribosomal proteins (MRPs) (Gerber et al., 2004; Lapointe et al., 2018). In glucose-replete conditions, Puf3 degrades mRNA of MRPs limiting mitochondrial translation (Houshmandi and Olivas, 2005; Olivas and Parker, 2000), which aligns with known metabolism preferences of yeast for glucose utilization through fermentation. In contrast, glucose depletion or switch to non-fermentable carbon sources leads to Puf3 phosphorylation and the subsequent switch of its function from mRNA degradation to the facilitation of translation (Lee and Tu, 2015). Our data indicate that expression of Cth1 and Cth2 is inversely correlated with the levels of mitochondrial ribosomal proteins during the transition of yeast cells from high glucose media to media containing glycerol further supporting the role of Cth1 and Cth2 in regulation of mitochondrial translation. Nonfermentable carbon sources, such as glycerol, require expression of the components of the oxidative phosphorylation and active mitochondrial translation (Couvillion *et al*., 2016), suggesting that the activity of Cth1/2 and Puf3 may allow proper synchronization between mitochondrial translation and Fe levels and that this regulation is not only involved in acute response to Fe deficiency, but may also play a role in physiological, homeostatic processes. Supporting this idea, we recently found that increased expression of Cth2 is limiting the expression of mitochondrial ribosomal proteins during yeast replicative aging (Patnaik et al., 2022).

The fact that genes encoding MRPs lack AREs in their 3’UTR suggests that Cth1 and Cth2 may regulate translation of mitochondrial ribosomal proteins indirectly by affecting phosphorylation of the Puf3 mRNA-binding protein. However, the nature of the signal that leads to phosphorylation of Puf3 and the specific kinases mediating the role of Cth1 and Cth2 in regulating mitochondrial translation warrants further investigation. Among potential candidates are PKA and Sch9, which have been shown to phosphorylate Puf3 when glucose becomes limited (Lee and Tu, 2015). In addition, phosphorylation of Puf3 in response to changes in the mitochondrial membrane potential has been recently reported to promote mitochondrial biogenesis in several mutants lacking components of the mitochondrial ETC (Liu et al., 2021).

Another effect of Fe deficiency on translation was identified when 3’UTR sequences were analyzed. Ribosome profiling of Fe-depleted cells revealed a striking decrease in the activity of the Fe-S cluster-containing Rli1 protein that serves as a ribosome recycling factor. We show that the number of footprints mapped to 3’UTR sequences was increased in Fe-deficient conditions to the same extent as seen previously in Rli1-depleted yeast cells (Young *et al*., 2015) leading to the generation of aberrant 3’UTR translation products. Notably, we observed a delayed response in the *cth1*Δ*cth2*Δ mutant indicating that Fe deficiency represses Rli1 function in wild-type cells as a result of Cth2-mediated post-transcriptional downregulation of *RLI1* transcripts. These data suggest that yeast cells deliberately downregulate the abundance of *RLI1* mRNA through Cth1 and Cth2 to limit Fe utilization by Rli1.

Finally, our study uncovered an important role of antisense lncRNA in the translational regulation of specific mRNA transcripts in response to low Fe. Our data demonstrate that Fe deficiency leads to increased expression of the regulatory antisense *MRS3* transcript repressing translation of cognate mRNA. Moreover, overexpression of the antisense lncRNA is sufficient to repress Mrs3 translation. This is, to the best of our knowledge, the first example of regulatory lncRNAs mediating the effect of Fe on protein translation. Recent genome-wide studies identified a number of genes that have overlapping antisense RNA transcripts in yeast and in other species (Liu et al., 2017; Till et al., 2018; Zhao et al., 2018) raising a possibility that expression of lncRNAs may have a role in Fe-dependent regulation of gene expression in higher eukaryotes. Additionally, regulation of protein translation by antisense transcripts has been implicated in several human diseases including cancer, cardiovascular and muscular pathologies, neurodegenerative disorders, and diabetes (Barman et al., 2019). It would be interesting to examine the importance of lncRNA-mediated translational repression in response to Fe deficiency in these pathophysiological processes.

Together, our findings uncovered a complex effect of Fe deficiency on the regulation of protein synthesis showing how this important metal affects protein translation at multiple levels. Given that many essential Fe-containing enzymes have been implicated in several steps of protein translation in mammals (Romero *et al*., 2021), the principles of Fe-dependent translational regulation uncovered in yeast may shed light on mechanisms that control protein synthesis in higher eukaryotes.

## Materials and Methods

### Yeast strains

The yeast strains used in this study and their genotypes are listed in Table S2. Prototroph W303a (*MATa, HIS3, TRP1, LEU2, URA3, ADE2, can1*) and W303a *cth1*Δ*cth2*Δ (*cth1*Δ*::hphB, cth2*Δ*::KanMX*) cells were cultured at 30°C in liquid SD medium (0.17% yeast nitrogen base without ammonium sulfate and without amino acids, 2% D-glucose, and 2 g/L Kaiser drop-out (Formedium)). For induction of Fe deficiency, cells were initially grown to OD_600_ = 0.2 and Fe^2+^ chelator bathophenanthrolinedisulfonic acid (BPS) was added to the final concentration 100 μM. Following the addition of BPS, cells were incubated for 3 h and 6 h. Cells were collected by rapid filtration through the 0.45 μm membrane filters using a glass holder filter assembly, scrapped with a spatula, and flash frozen in liquid nitrogen. Yeast strains encoding HA-tagged Mrs3 and Dph1 proteins were generated using CRISPR/Cas9 genome editing as described (Barre et al., 2020).

### Ribo-Seq and RNA-seq sequencing, data processing, and analysis

Yeast extracts were prepared by cryogrinding the cell paste with BioSpec cryomill. The cell paste was re-suspended in lysis buffer (20 mM Tris-HCl pH 8, 140 mM KCl, 5mM MgCl_2_, 0.5 mM DTT, 1% Triton X-100, 100 μg/mL cycloheximide), spun at 20,000 x g at 4°C for 5 min and 1 mL of the supernatant was divided into two tubes, for total mRNAs and footprint extraction. For Ribo-Seq, 50 OD_260_ units of the lysate (in 1 mL of lysis buffer) were treated with 10 μL of RNase I (100 U/μL) for 1 h, with gentle rotation of samples at room temperature. Then, 1 mL of RNase-treated lysate was layered on a sucrose gradient for isolation of monosomes using ultracentrifugation followed by footprint extraction with hot acid phenol method. For RNA-Seq samples, poly(A) mRNA isolation was performed using a poly(A) mRNA isolation kit with subsequent mRNA fragmentation. The RNA-seq and Ribo-Seq libraries were prepared as described previously (Beaupere et al., 2017) and sequenced using the Illumina HiSeq platform.

For Ribo-Seq data, the adapter sequence was removed using Cutadapt 4.1 (Martin, 2011) and reads less than 23 nucleotides were filtered out. The Ribo-Seq and RNA-Seq reads were aligned to the *S. cerevisiae* genome from the Saccharomyces Genome Database (https://www.yeastgenome.org/, release number R64-2-1). Sequence alignment was performed using STAR software 2.7.1a, allowing two mismatches per read (Dobin et al., 2013). Counts were generated with featureCounts from Rsubread 1.22.2 package (Liao et al., 2019). We filtered out genes with low number of reads (less than 10 counts in less than 66.6% of samples) resulting in 5506 detected genes in RNA-Seq and 5069 genes in Ribo-Seq expression matrices. Identification of differentially expressed genes was performed using the generalized linear model of the EdgeR package (GLM, glmFit, glmLRT) with an adjusted p-value cutoff (FDR<0.05) (Robinson et al., 2010).

### Ribosome occupancy analysis

To estimate translational efficiency changes during Fe deficiency, we analyzed ribosomal occupancy (RO), which represents a ratio between ribosomal footprints and mRNA abundance allowing to identify actively translated transcripts. To calculate RO, genes with low counts (less than 10 counts) were filtered out from RNA-Seq and Ribo-Seq datasets containing raw counts. RNA-Seq and Ribo-Seq data were then RLE normalized (“Relative Log Expression” normalization) using the edgeR package (Robinson *et al*., 2010) and RO was calculated using the following formula: log2 (Ribo-seq counts +1) – log2 (RNA-seq counts + 1). We used Limma R package to estimate the contribution of transcriptional and translational regulation to gene expression changes and identify translationally regulated genes (Anders and Huber, 2010; Ritchie et al., 2015).

### Quantification of antisense transcripts

To quantify antisense transcripts, sequence reads that align to annotated ORFs and are transcribed from the opposite DNA strand were counted using featureCounts from Rsubread 1.22.2 package (Liao *et al*., 2019). First, we filtered out genes with less than 5 counts in less than 66.6% of samples (an empirically chosen threshold aiming to bring the distribution of the counts closer to normal and preserve the maximum number of genes with minimal variation between replicates). After filtering step, we obtained 726 genes. RLE normalized counts were then used to identify differentially expressed antisense transcripts during Fe deficiency using Limma R package.

### Polysome analysis

Polysome analysis was performed according to the previously published protocol (Ramos-Alonso *et al*., 2018). Cells were grown in SC media overnight upon 0.2 OD_600_ density and supplemented with 100 μM of the bathophenanthroline disulfonate (BPS) to induce Fe deficiency. Cells were collected after 3 and 6 hours of treatment and resuspended in 700 _µ_l of lysis buffer [20 mM Tris-HCl, pH 8, 140 mM KCl, 5 mM MgCl2, 0.5 mM dithiothreitol, 1% Triton X-100, 0.1 mg/mL CHX, and 0.5 mg/mL heparin]. Aliquots of cell extracts containing 8.5 OD260 units were loaded on top of sucrose gradients (5-50% wt/wt). The gradients were sedimented at 35,000 rpm at 4°C in a SW41 rotor (Beckman) for 2 h 40 min. Fractions were analyzed by UV detection at 260 nm.

### RT-qPCR

Total RNA was isolated by hot acid phenol extraction. RNA was treated with DNaseI, and 1 μg of RNA was used for cDNA synthesis using SuperScript III reverse transcriptase (Thermo Fisher Scientific) with random hexamer primers according to manufacturer’s instructions. mRNA expression was then analyzed by real-time PCR using KAPA SYBR FAST qPCR Master Mix (Kapa Biosystems) and the CFX-96 Touch Real-Time PCR Detection System (Bio-Rad Laboratories). *ACT1* was used as a reference gene for normalization of mRNA expression between genotypes. The primers used for RT-qPCR are listed in Table S2. Results are represented as means ± SEM from three independent experiments.

### Western blot analysis

Total protein extracts were prepared using TCA precipitation. 5 OD_600_ units of cells were resuspended in 0.5 mL of 6% TCA, incubated at least 10 min on ice and centrifuged for 5 min at 4°C at maximum speed (∼14,000 x g). Next, the pellet was washed twice with acetone and air-dried. The pellet was then resuspended in 250 μL of HU buffer (5 M urea, 50 mM Tris-HCl pH 7.5, 1% SDS, 1 mM PMSF) and samples were homogenized with glass beads by vortexing at maximum speed for 10 x 30 sec. Equal amounts of proteins were resolved in 10% SDS-PAGE gels and transferred to PVDF membrane. Anti-HA-Peroxidase, High Affinity (3F10) rat monoclonal antibody (Roche) was used to detect HA-tagged proteins. Mouse anti-Pgk1 monoclonal antibody (Life Technologies) and HRP-conjugated secondary anti-mouse antibody (Santa Cruz Biotechnology) were used to detect Pgk1 protein levels as a loading control.

### Expression of the *MRS3^AS^* lncRNA

To generate plasmids that express the *MRS3^AS^*lncRNA, we cloned *MRS3* ORF (or indicated fragments) in reverse orientation under the control of *TEF1* promoter into pCEV plasmid (Vickers et al., 2013) containing Zeocin (ZeoMX) resistance. *W303a* yeast cells were transformed with *pCEV-MRS3^AS^* plasmids and selected on media containing Zeocin at a final concentration of 300 μg/mL. Expression of the antisense *MRS3^AS^* transcript was verified using RT-qPCR.

### Quantification and statistical analysis

Statistical analysis was performed using Prism 9.3.1 (GraphPad Software, Inc). Statistical significance of the RT-qPCR data was determined by calculating p values using one-way ANOVA. Error bars represent standard errors of the means (SEM). The information about number of replicates and P values can be found in figure legends. For RNA-Seq and Ribo-Seq analyses tree biological replicates were analyzed per each condition.

## Data availability

Raw reads and processed sequencing data generated by this study have been deposited in the GEO database (www.ncbi.nlm.nih.gov/geo/; Accession: GSE193025)

## Acknowledgments

We would like to thank Dr. Alan Hinnebusch for providing yeast strains containing 3xHA tags in the 3’UTRs of *SED1* and *CWP2* used in this study. This work was supported by NIH Grants AG058713 and AG066704 to VML, and grants PID2020-116940RB-I00 and CEX2021-001189-S from MCIN/AEI/10.13039/501100011033 to SP.

## Competing interests

The authors declare no competing interests.

## Supplementary Figures

**Fig. S1.**
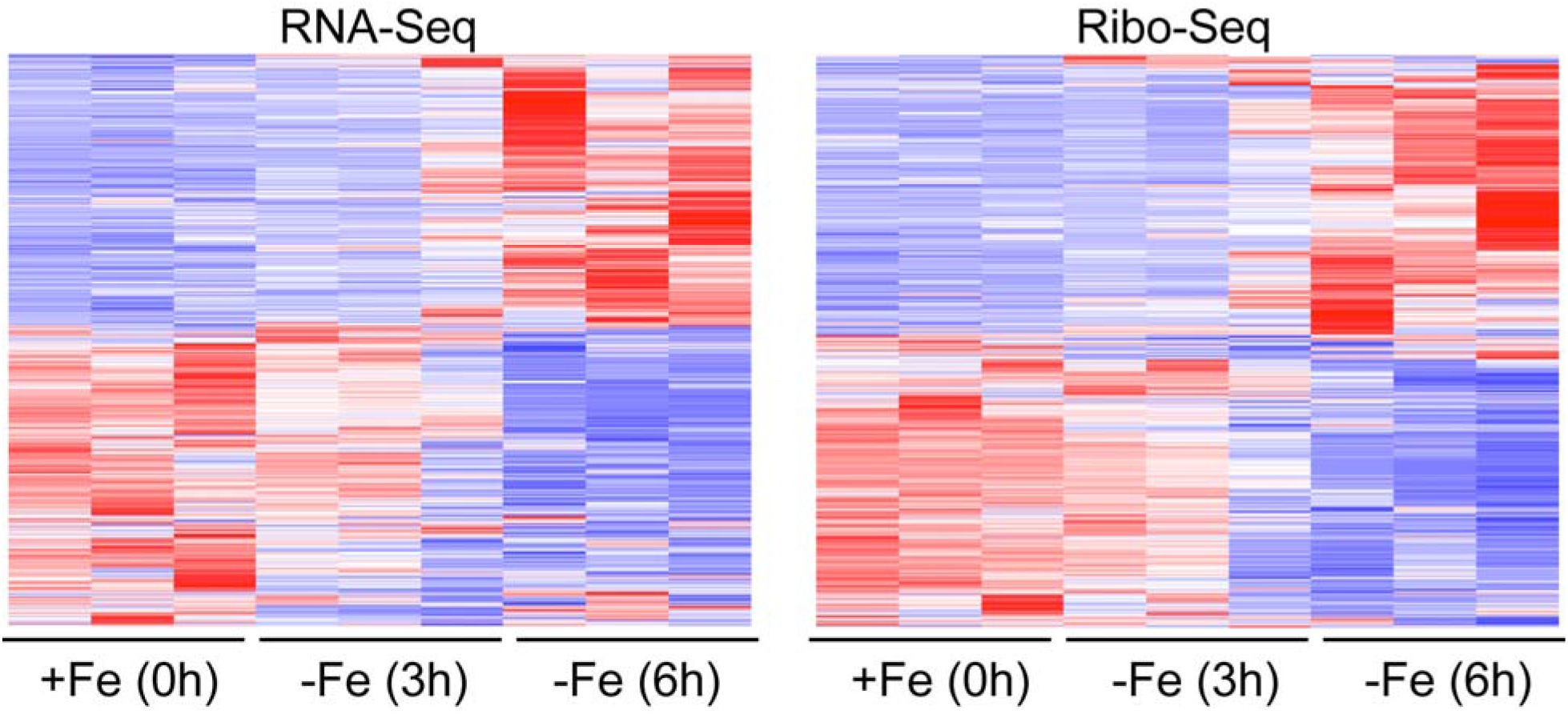
Changes in RNA-Seq and Ribo-Seq expression profiles in response to Fe deficiency. Heatmaps show changes in transcription (RNA-Seq) and protein translation (Ribo-Seq) scaled expression values (RLE-normalized log2 read counts) in wild-type cells during short-term (3h) and prolonged (6h) Fe deficiency.

**Fig. S2.**
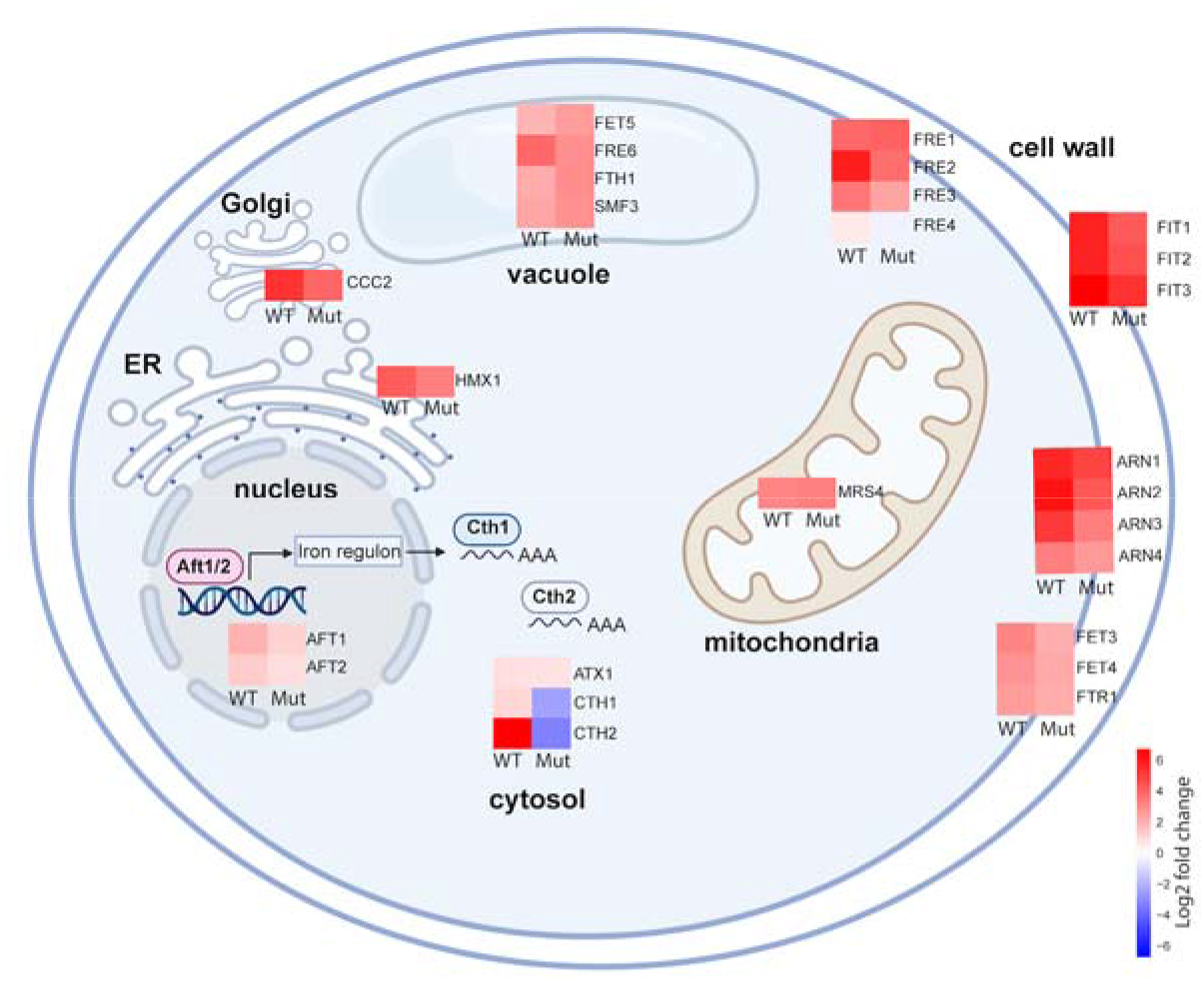
Heatmaps show log2 fold change in protein translation (Ribo-Seq) of Fe regulon genes during prolonged (6h) Fe deficiency in wild-type (WT) cells and the *cth1*Δ*cth2*Δ mutant (Mut) (FDR<0.05).

**Fig S3.**
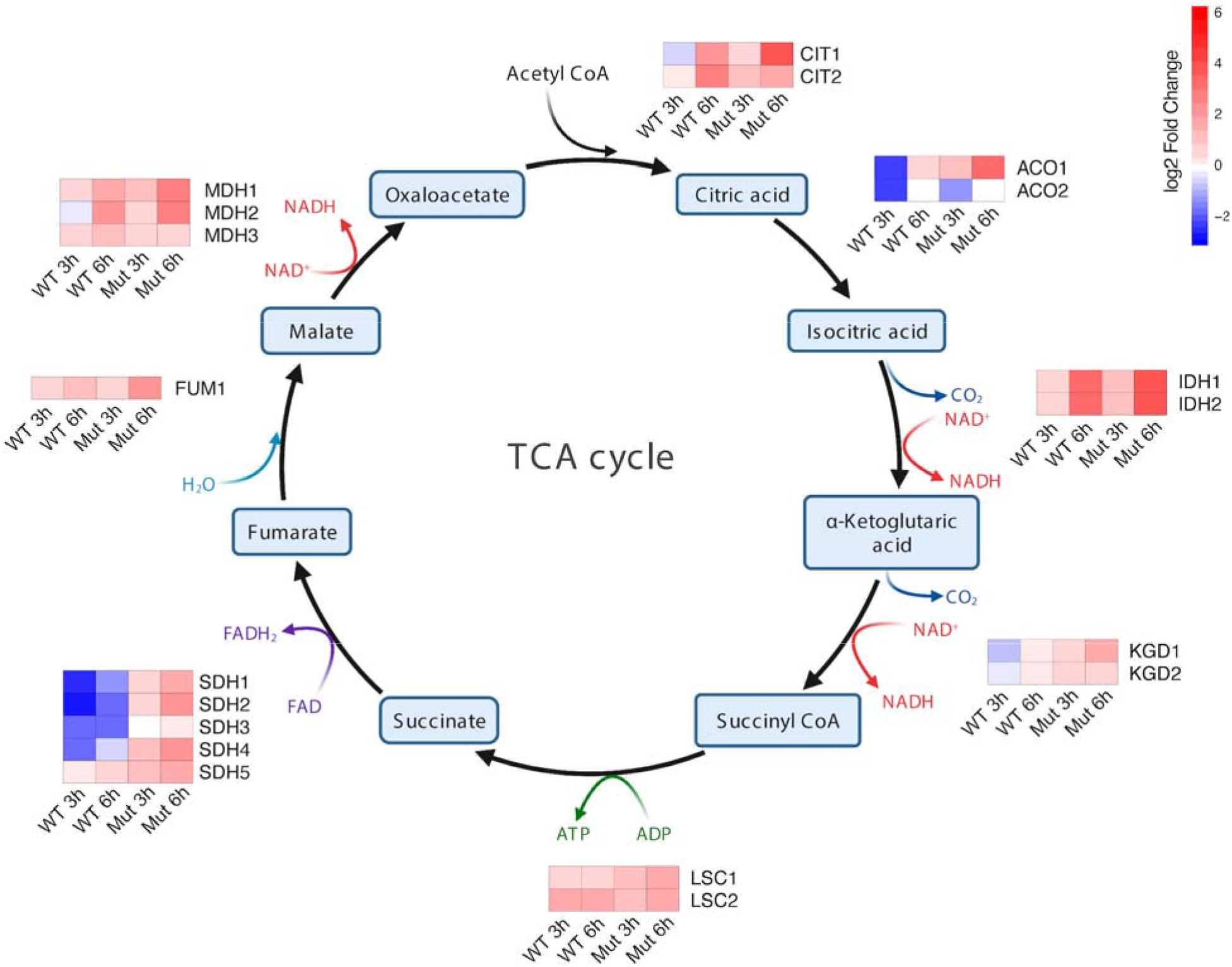
Heatmaps show log2 fold change in protein translation (Ribo-Seq) of differentially regulated genes of the TCA cycle during short-term (3h) and prolonged (6h) Fe deficiency in wild-type (WT) cells and the *cth1*Δ*cth2*Δ mutant (Mut) (FDR<0.05).

**Fig S4.**
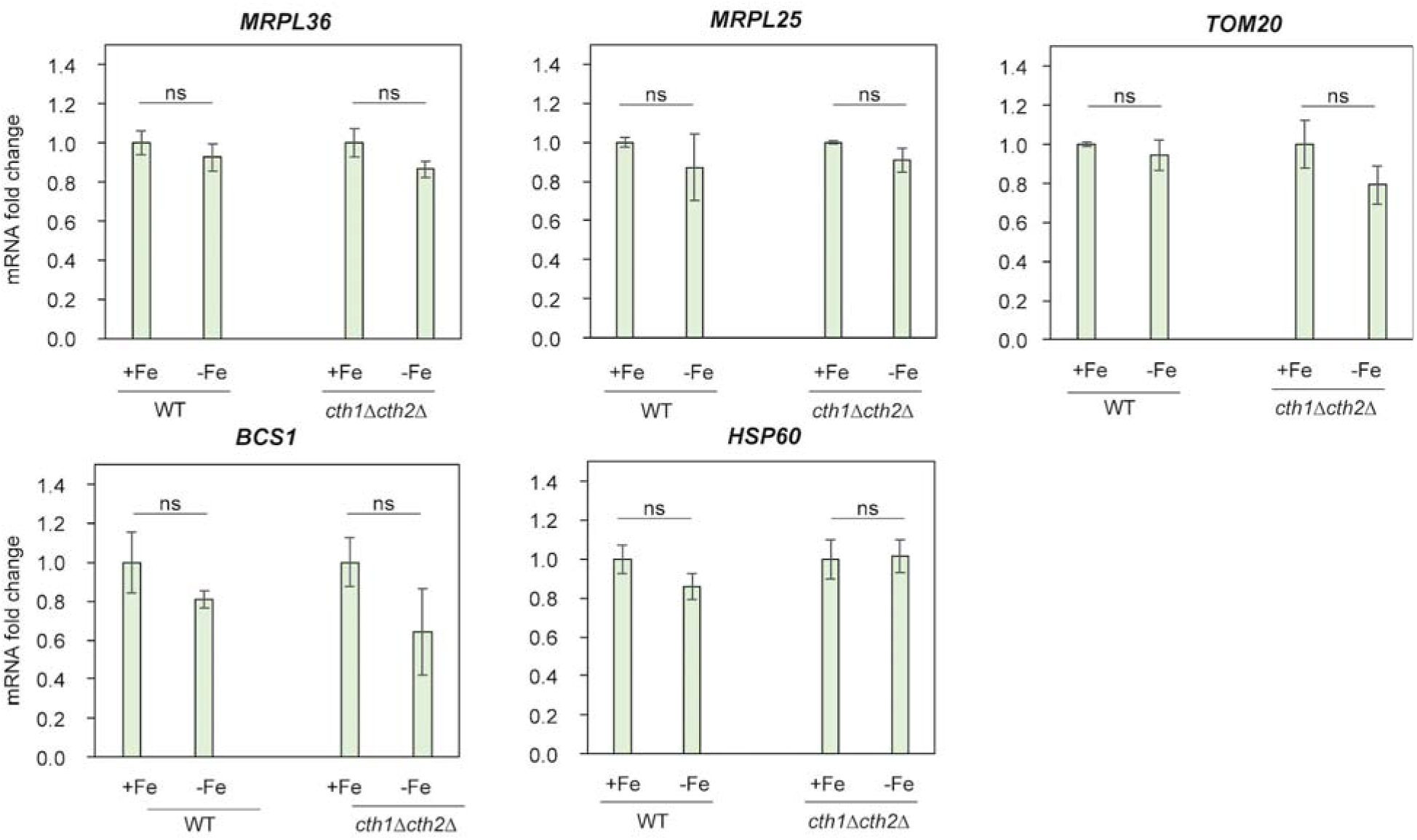
Fe deficiency does not affect transcription levels of mitochondrial ribosomal proteins. The expression of *MRPL36, MRPL25, TOM20, BCS1,* and *HSP60* was determined by RT-qPCR. Error bars represent SEM of three independent experiments; ns, non-significant (one-way ANOVA).

**Fig S5.**
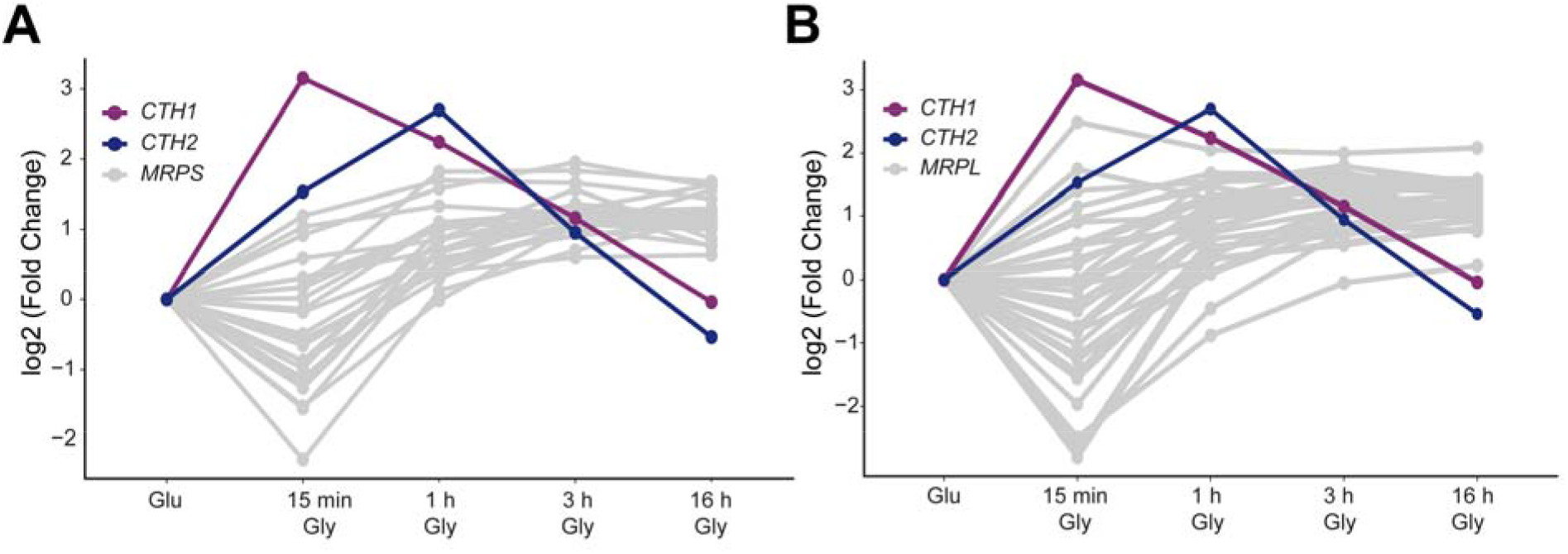
Expression of the mitochondrial ribosomal proteins of the small subunit (A) and large subunit (B) during the shift from the fermentable to non-fermentable carbon source. Cells were grown in YPD (Glu) medium to log phase and shifted to YPG (Gly) medium for the indicated time. Fold change in protein translation was measured by Ribo-Seq.

**Fig S6.**
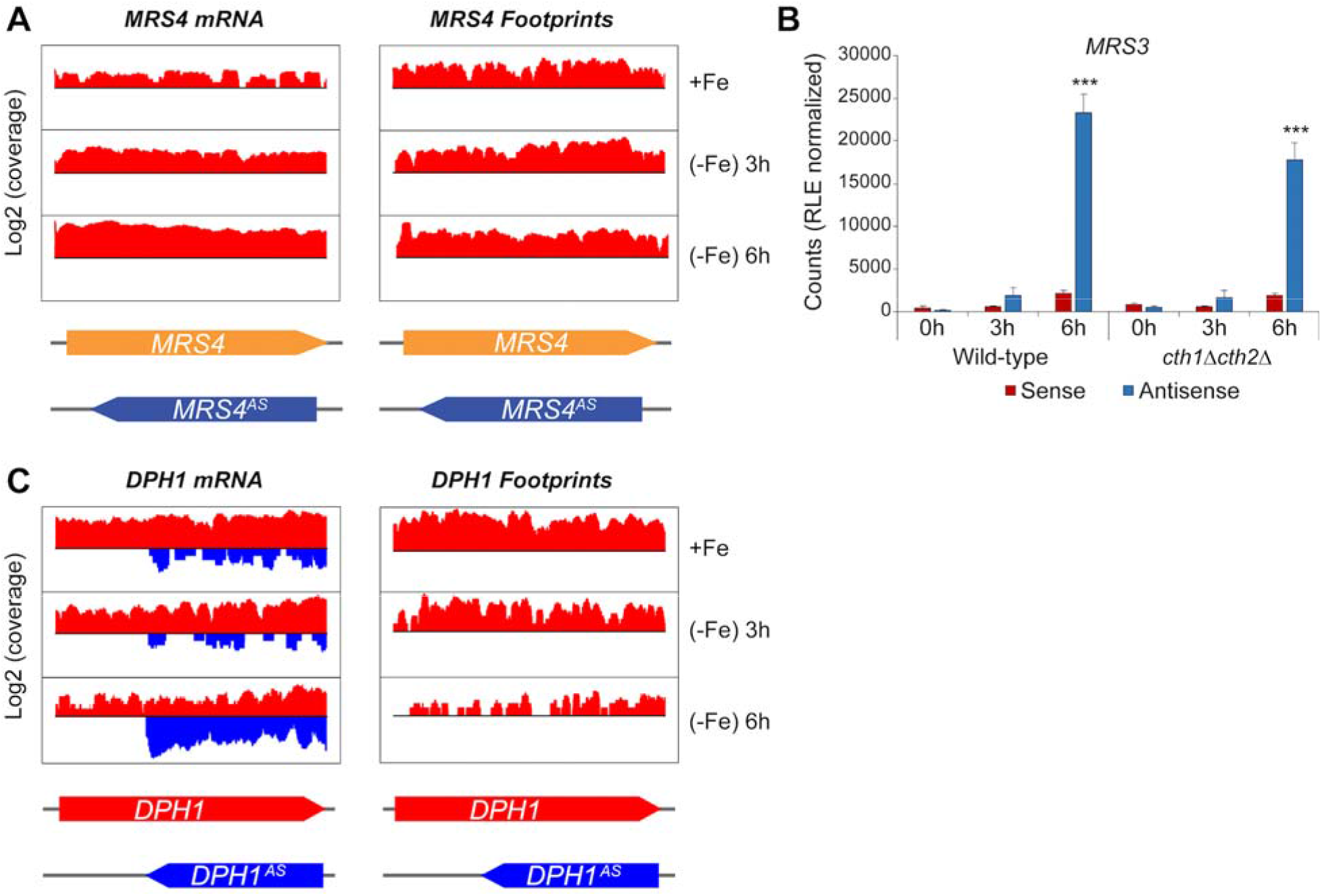
Expression of antisense lncRNA is a conserved regulatory mechanism in yeast. **A**) Coverage plot of mRNA and footprint reads mapped to sense (red) and antisense (blue) *MRS4* transcript. **B)** Expression of antisense *MRS3* (*MRS3*^AS^) transcript during Fe deficiency is independent of Cth1 and Cth2. Relative expression of *MRS3* and *MRS3*^AS^ transcripts was analyzed by RT-qPCR. Error bars represent SEM of three independent experiments, ***p<0.001 (two-way ANOVA). **C**) Fe deficiency induces expression of antisense *DPH1* (*DPH1*^AS^) transcript. Figure shows coverage plots of mRNA and footprint reads mapped to sense (red) and antisense (blue) *DPH1* transcript.

**Table S1.**
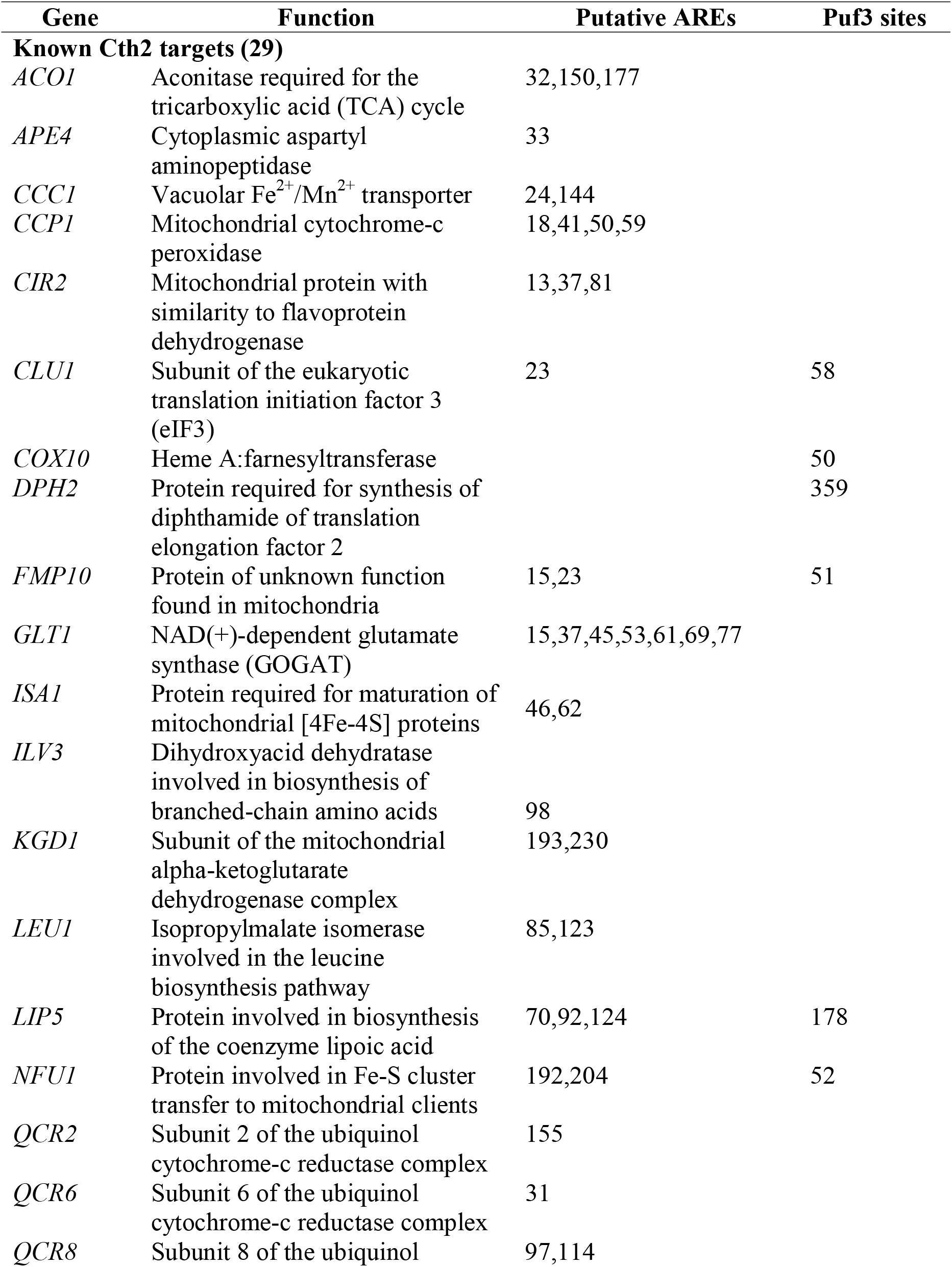
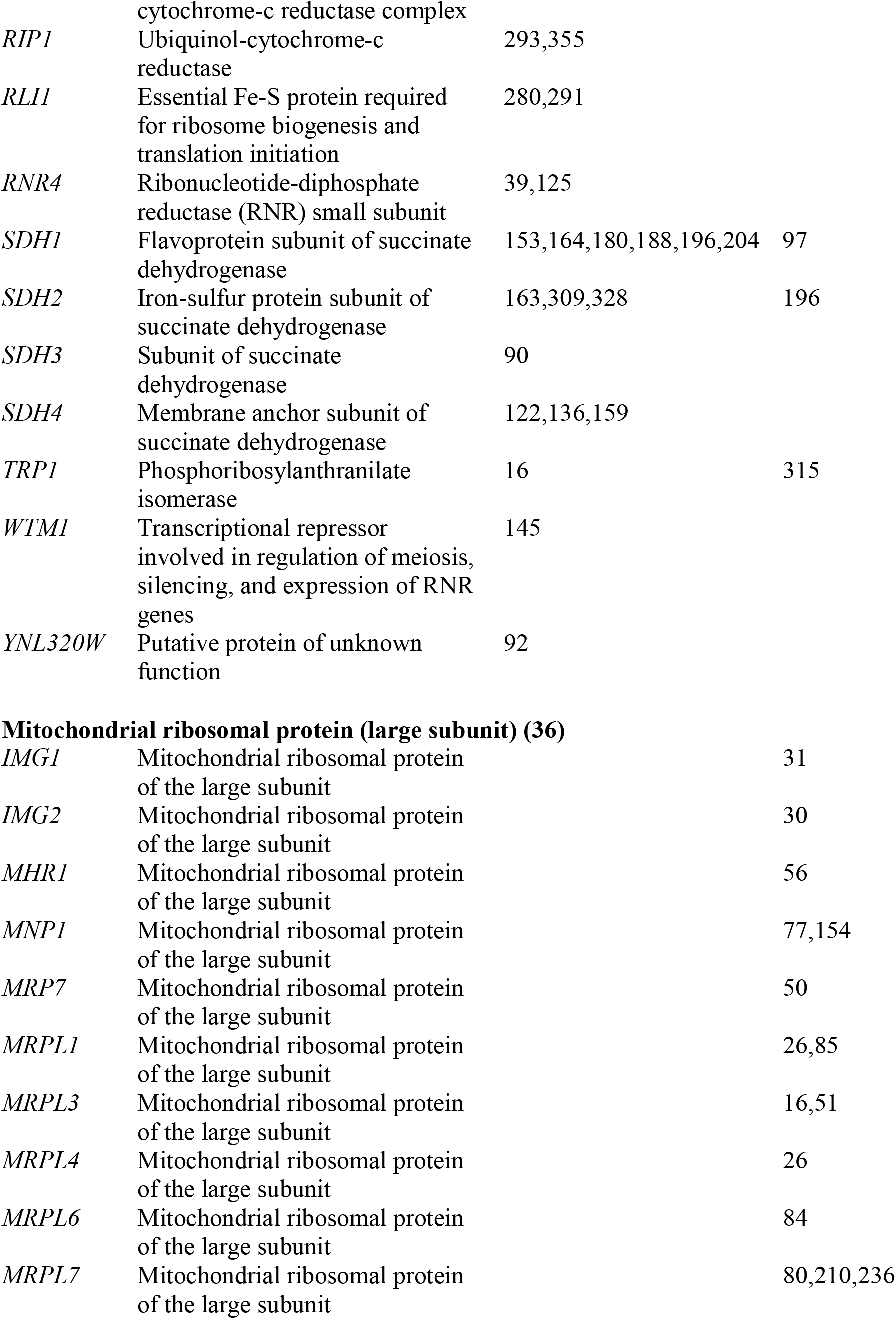
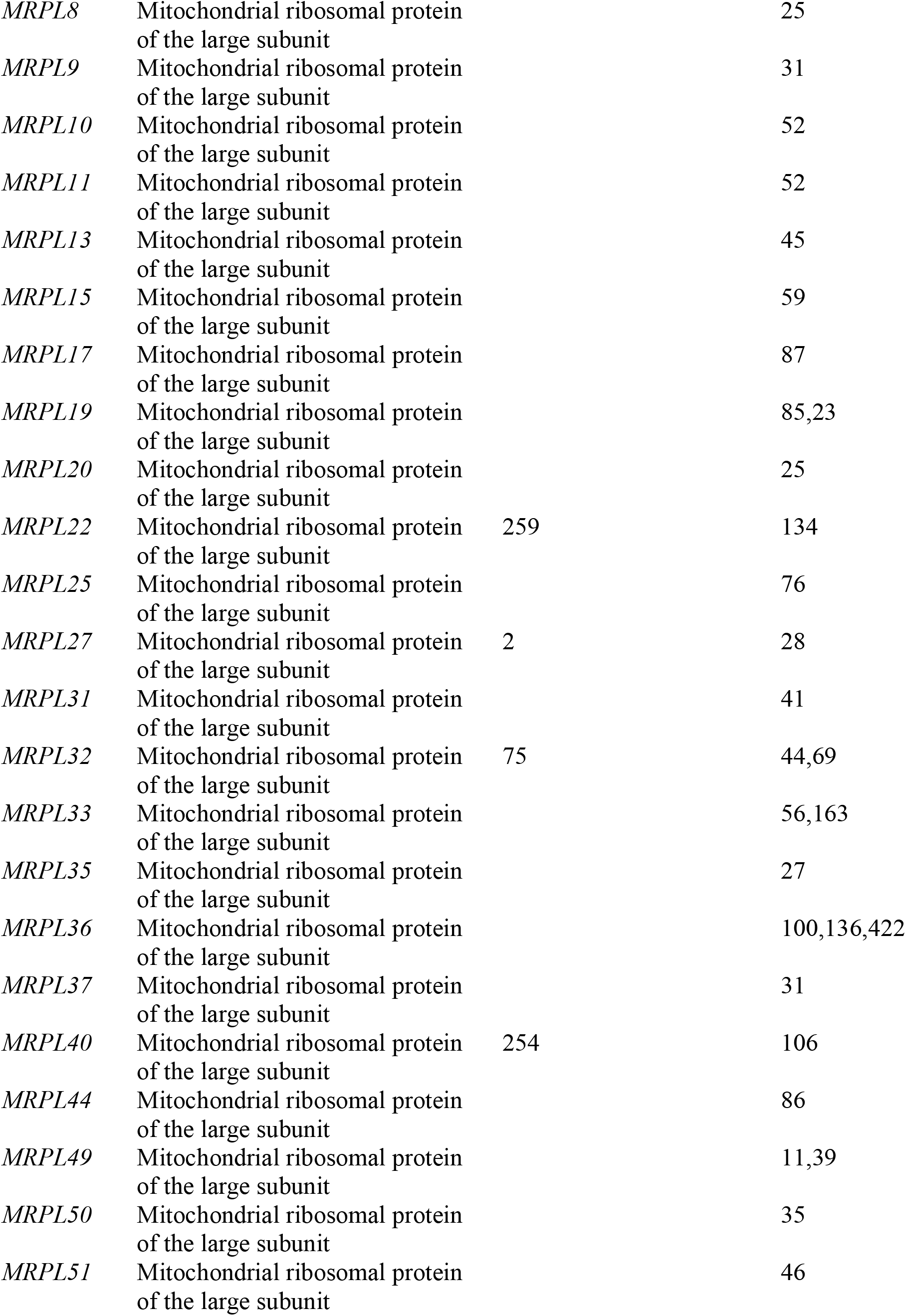
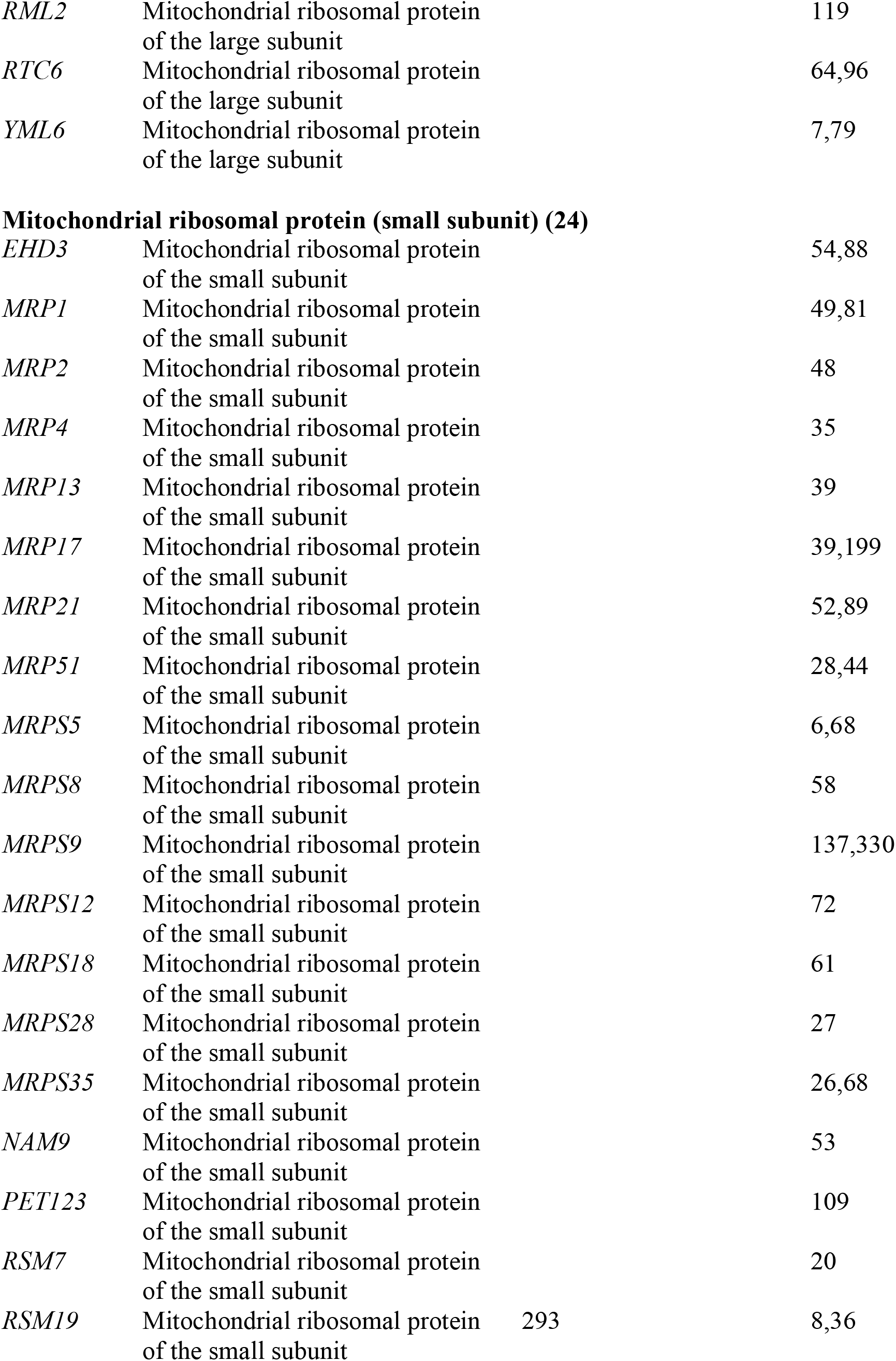
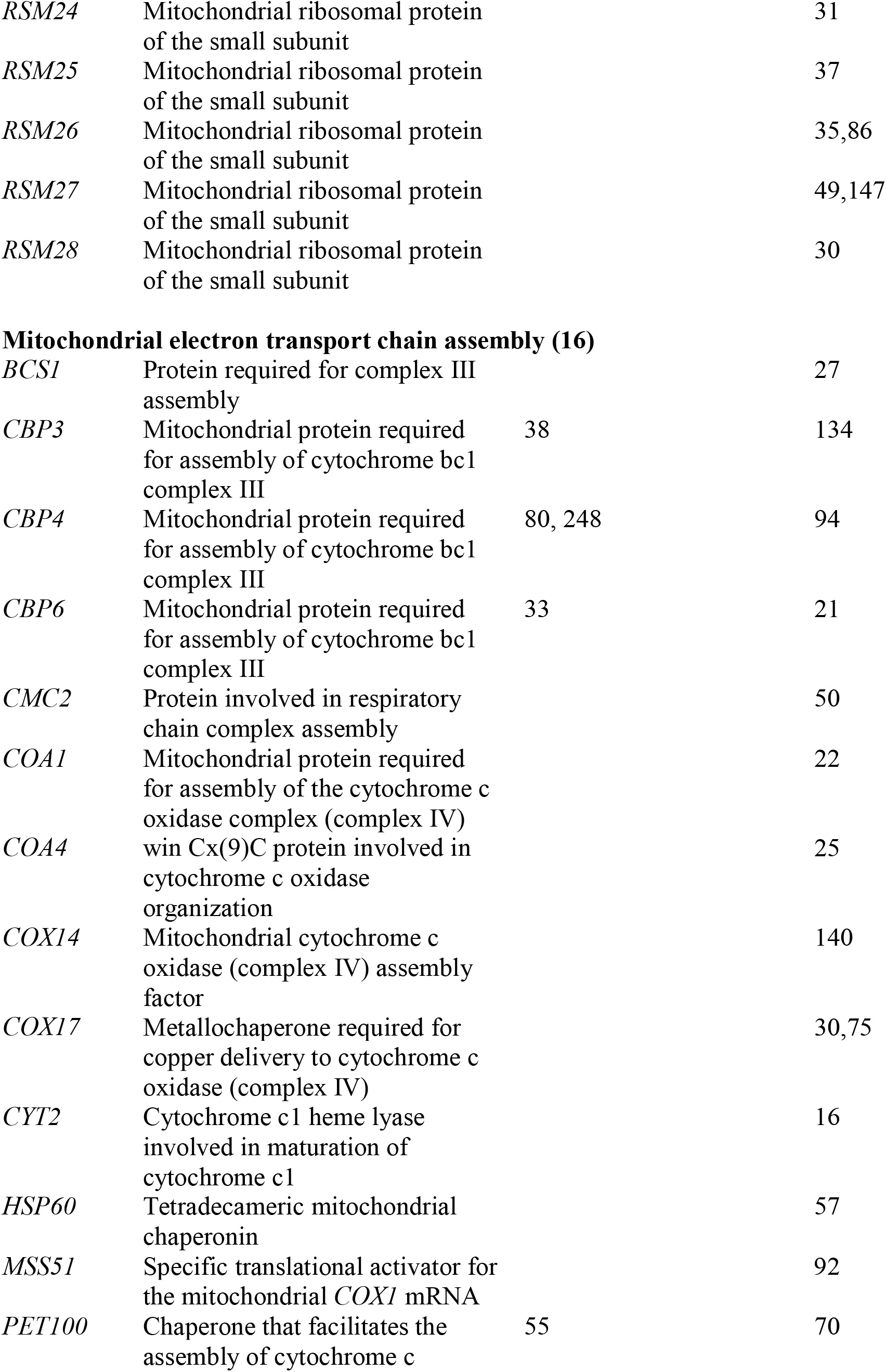
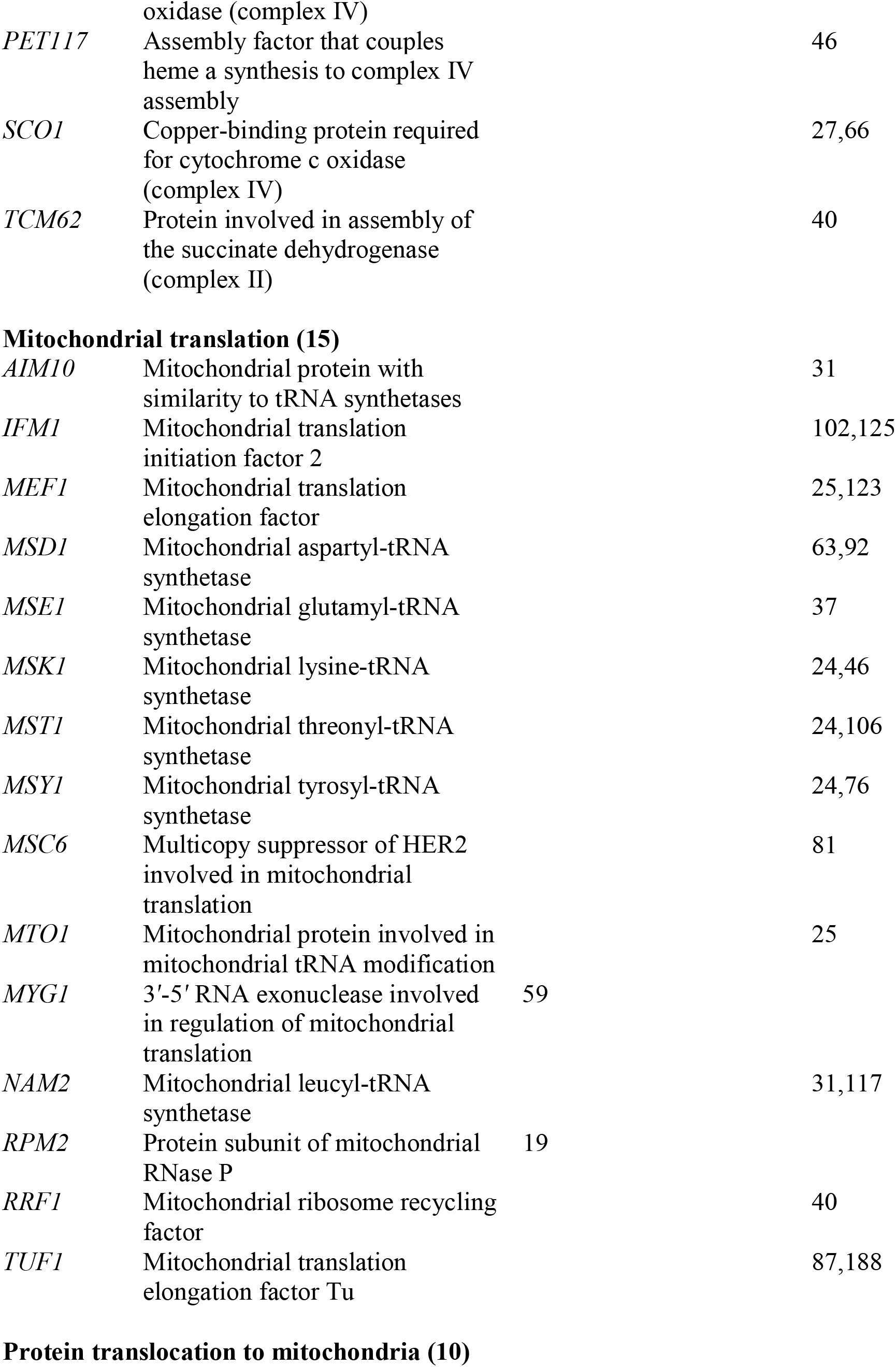
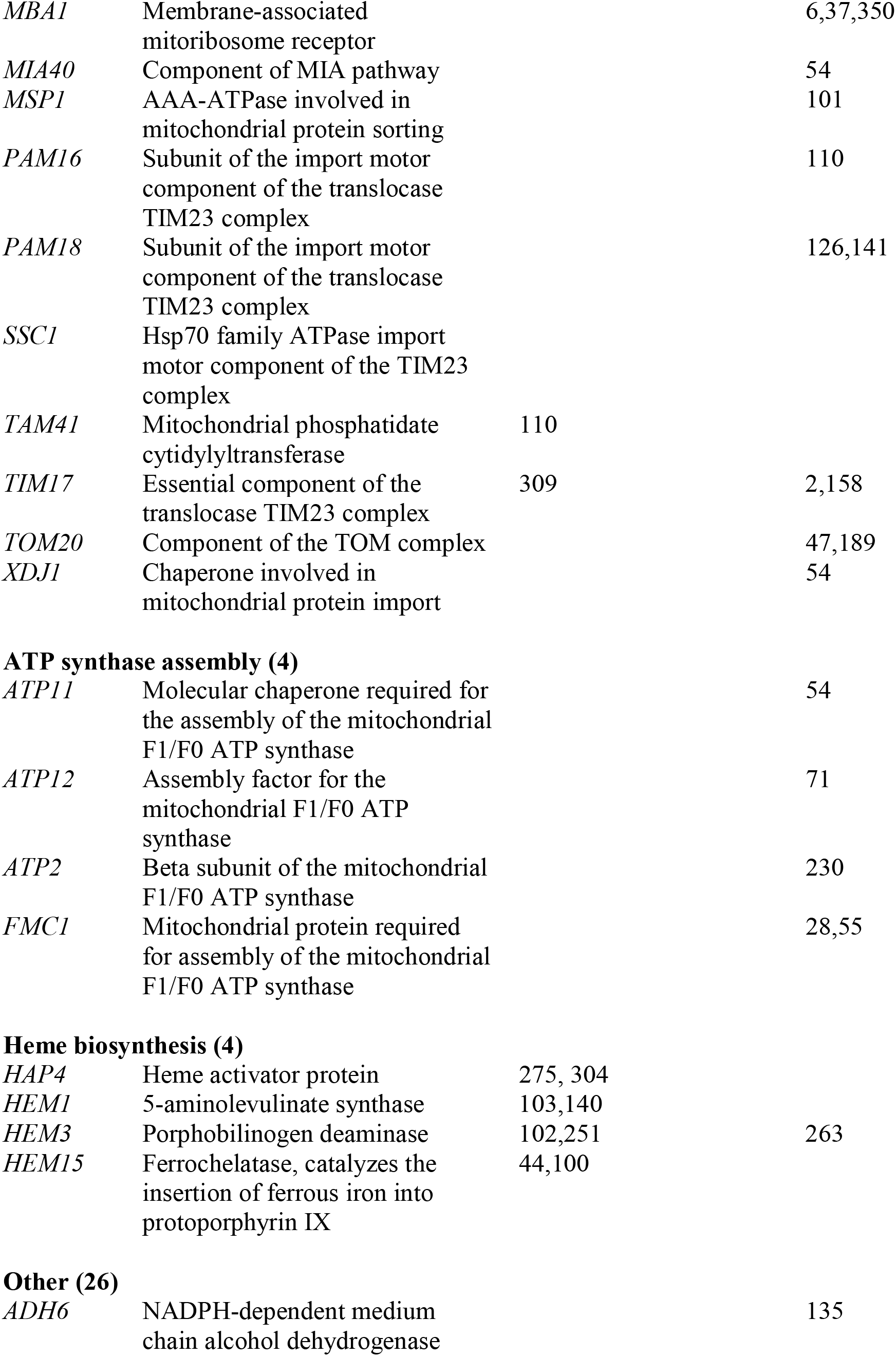
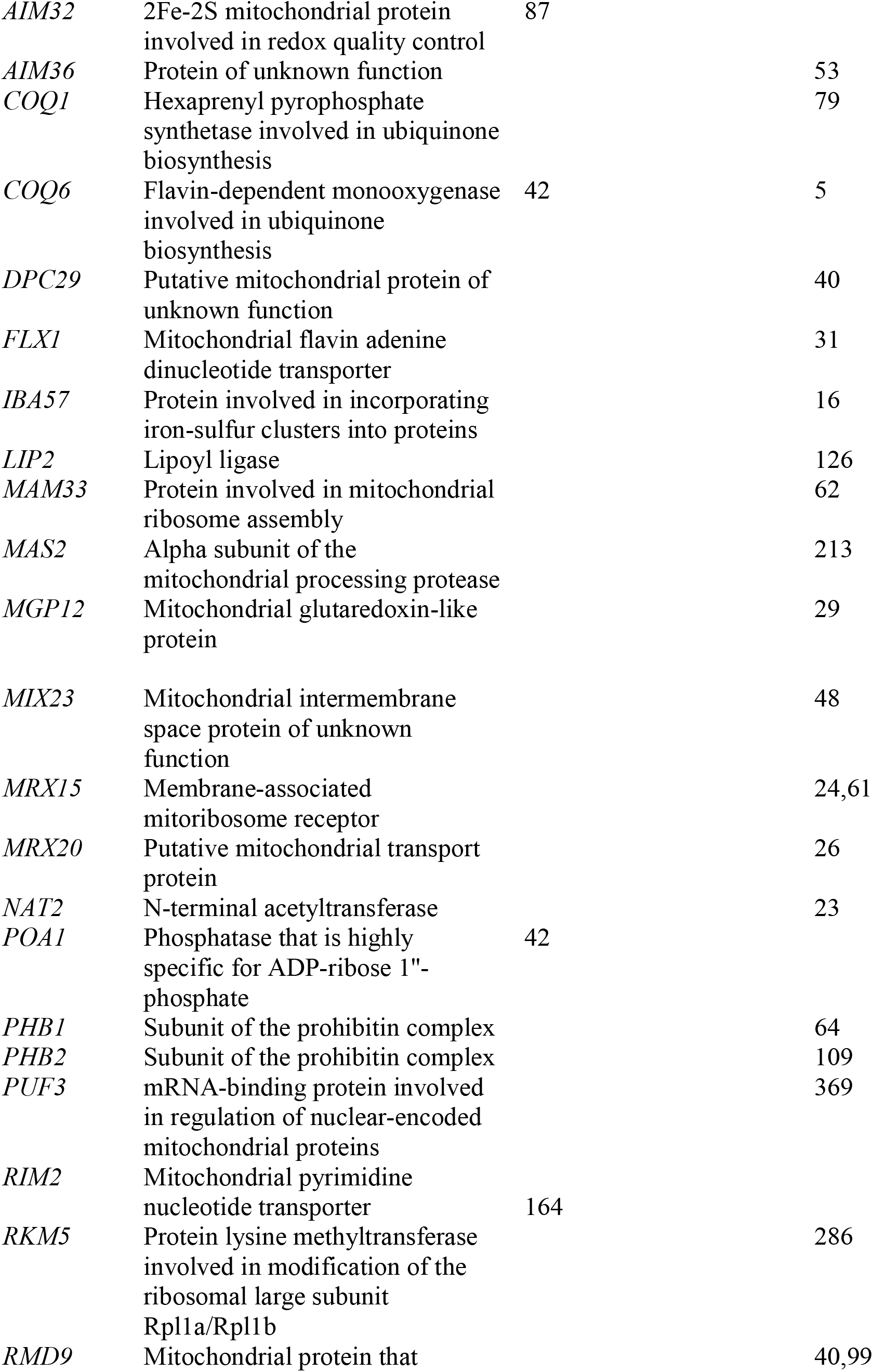
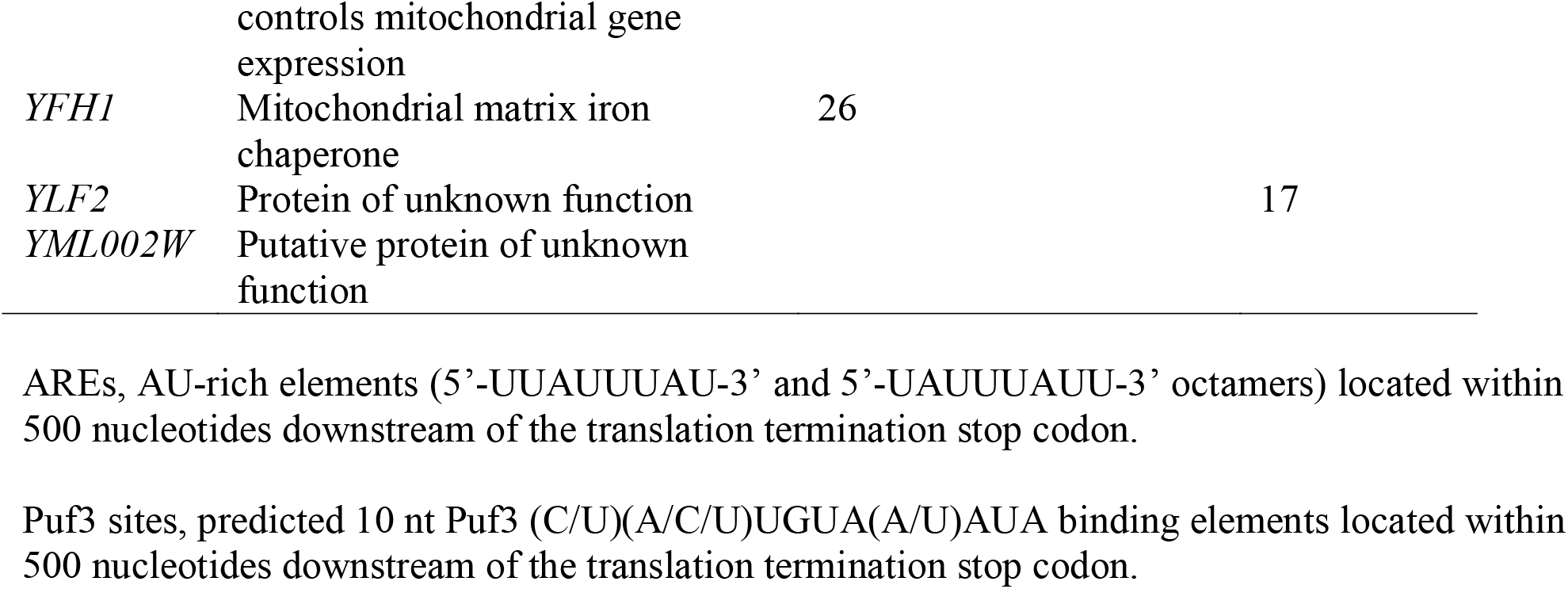
Location of putative AU-rich elements (AREs) and Puf3 binding sites in genes translationally regulated by yeast Cth1/Cth2 proteins in response to Fe deficiency.

**Table S2.**
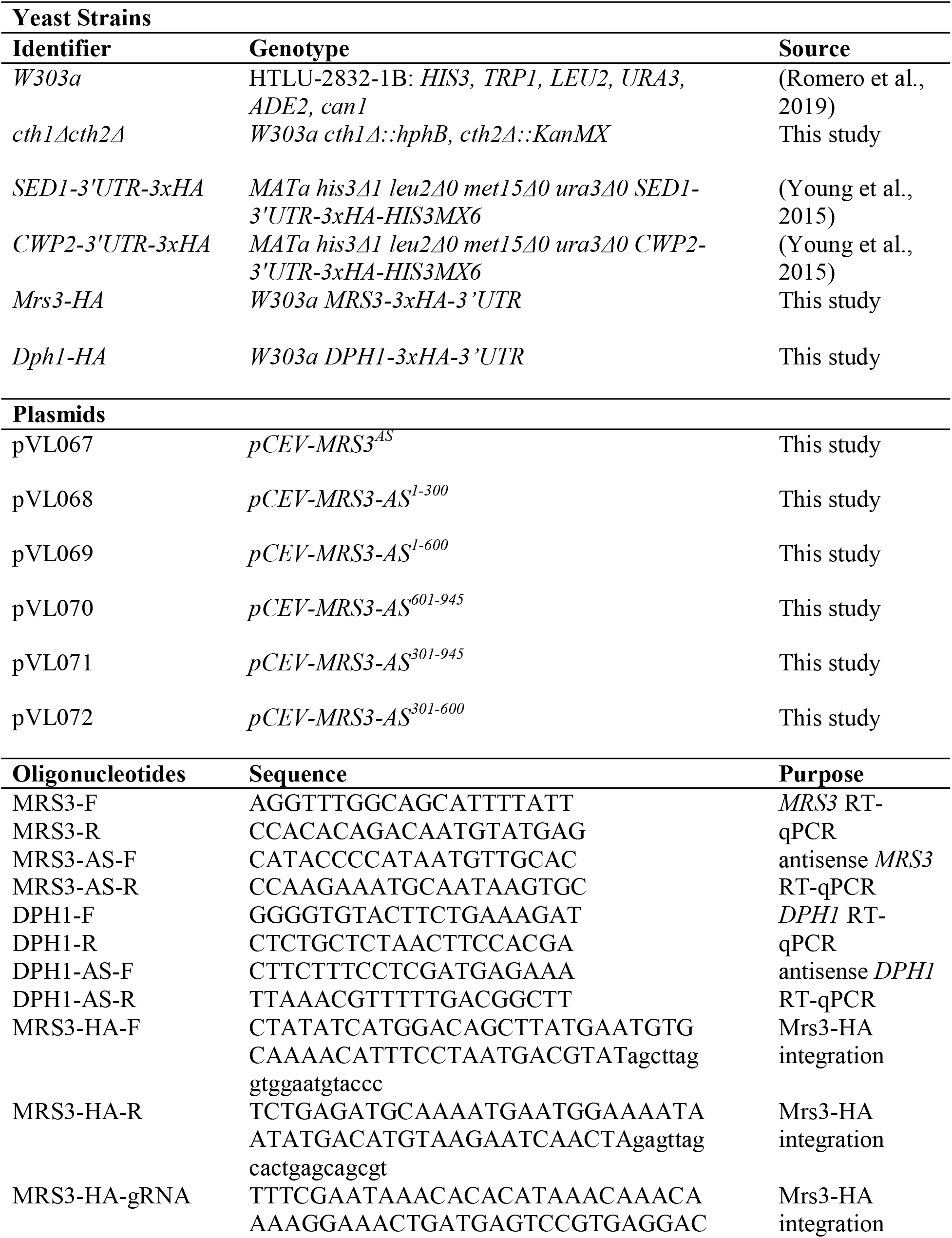
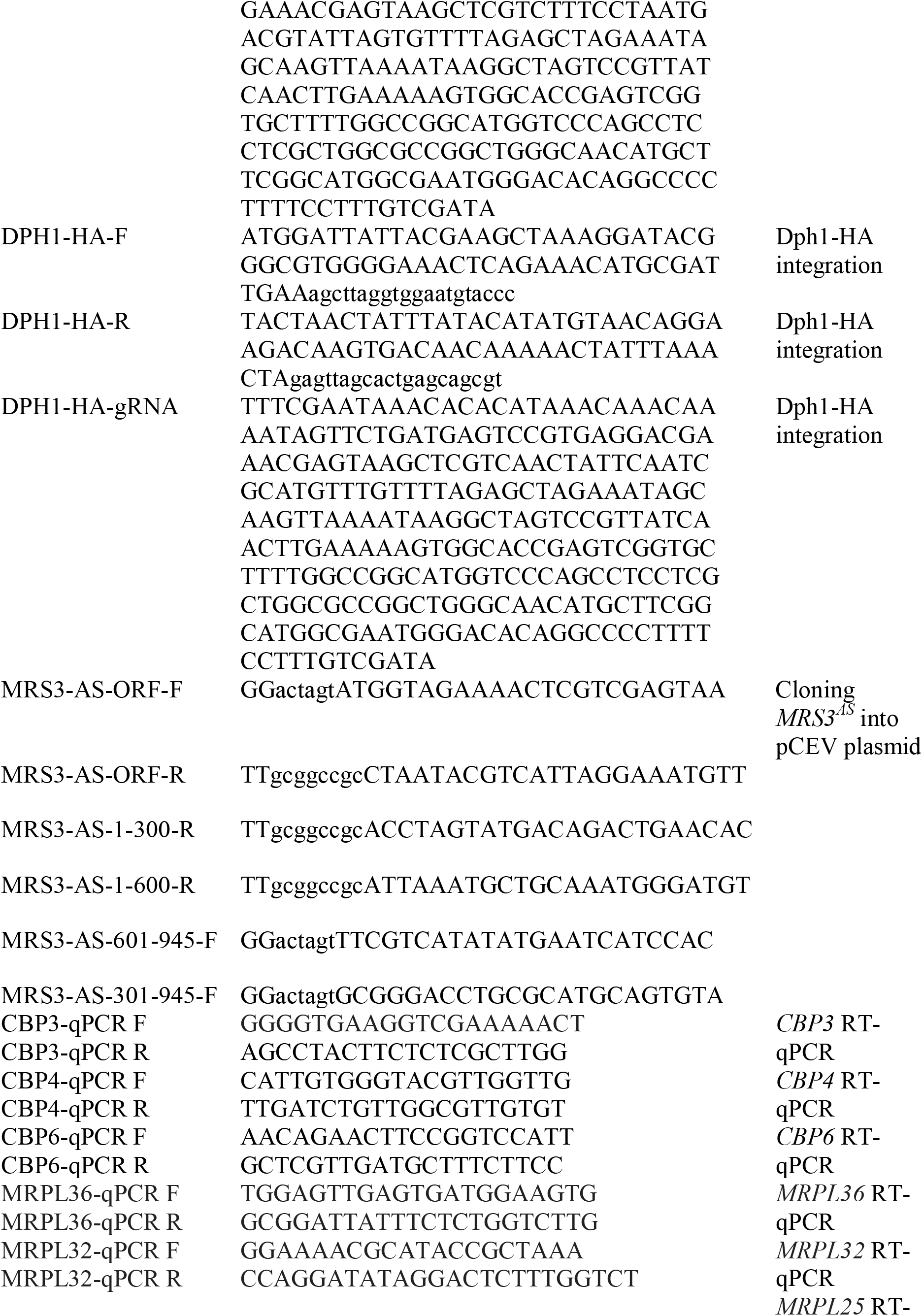
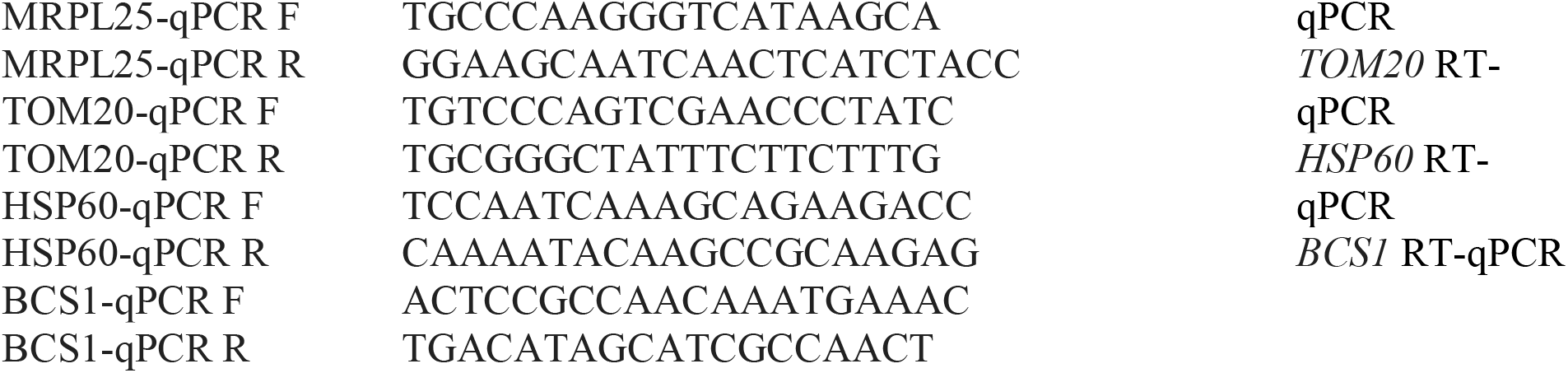
Key resources table.

## Notes

### Competing Interest Statement

The authors have declared no competing interest.

### Summary of Updates

This version of the manuscript includes additional data showing that iron deficiency influences the translation of genes involved in heme biosynthesis (Revised Figure 2, Panel F) and regulates the transcription of regulatory lncRNAs (new Figure 4).

